# Exploration and analysis of molecularly annotated, 3D models of breast cancer at single-cell resolution using virtual reality

**DOI:** 10.1101/2021.06.28.448342

**Authors:** Dario Bressan, Claire M. Mulvey, Fatime Qosaj, Robert Becker, Flaminia Grimaldi, Suvi Coffey, Sara Lisa Vogl, Laura Kuett, Raul Catena, Ali Dariush, Carlos Gonzalez-Fernandez, Eduardo A. Gonzalez-Solares, Mohammad Al Sa’d, Aybüke Küpcü Yoldaş, Tristan Whitmarsh, Ilaria Falciatori, Spencer S. Watson, CRUK IMAXT Grand Challenge Team, Johanna A. Joyce, Nicholas Walton, Bernd Bodenmiller, Owen Harris, Gregory J. Hannon

**Author notes:** These authors contributed equally (Dario Bressan, Claire M. Mulvey, Fatime Qosaj).

## Abstract

A set of increasingly powerful approaches are enabling spatially resolved measurements of growing numbers of molecular features in biological samples. While important insights can be derived from the two-dimensional data that many of these technologies generate, it is clear that extending these approaches into the third and fourth dimensions will magnify their impact. Realizing biological insights from datasets where thousands to millions of cells are annotated with tens to hundreds of parameters in space will require the development of new computational and visualization strategies. Here, we describe Theia, a virtual reality-based platform, which enables exploration and analysis of either volumetric or segmented, molecularly-annotated, three-dimensional datasets, with the option to extend the analysis to time-series data. We also describe our pipeline for generating annotated 3D models of breast cancer and supply several datasets to enable users to explore the utility of Theia for understanding cancer biology in three dimensions.

## INTRODUCTION

The ability to interrogate tissues at the single cell level and to gather genomic and proteomic information at scale for the individual constituents of living systems is having an enormous impact. Single-cell RNA and DNA sequencing is revealing new cell types, enabling comprehensive catalogues of human and mouse tissues, and revealing genetic and phenotypic heterogeneity in tumours. Yet, most normal and disease development happens in a spatial context. The three-dimensional architecture of a tissue has profound impacts on its function, whether it be through the establishment of morphogen gradients in development or through the effect of local tissue niches featuring distinct cell compositions or signalling states. However powerful, measurements made on disaggregated cells or nuclei generally lack this critical contextual information. In cancer, there is a growing appreciation that the initiation of disease, its progression toward invasion and metastasis, and its response to therapy are all profoundly influenced not only by the properties of the tumour cells themselves but also by the characteristics of the tumour microenvironment (TME)^1–3^. A description of the relationships between a tumour and its TME requires not only an enumerated catalogue of its constituent cell types and clonal lineages but also an understanding of their interactions and arrangement in three-dimensional space.

The desire to understand development, tissue architecture, and disease in context has prompted a drive toward the development of methods for making multiplexed measurements on intact tissues or tissue sections. Such methods measure portions of the transcriptome or proteome or alternatively can quantify metabolites. Some are destructive, while others leave tissue intact with the potential for layering additional measurements, and though a few methods are compatible with multimodal measurements, generally the number of markers measured outside of the primary modality is rather low. MERFISH^4, 5^, STARmap^6^, seqFISH^7^, Spatial Transcriptomics^8^, and SlideSeq^9^ can interrogate the expression of 100s to >10,000 transcripts *in situ*, with most of these approaches operating at cellular or sub-cellular resolution. Imaging Mass Cytometry (IMC)^10^, CODEX^11^, CycIF^12^, 4i^13^, and others can measure the expression of dozens of proteins simultaneously or serially. All of these are now beginning to produce new biological insights into the organization of tissues in two-dimensions. For example, recent studies using IMC have revealed that the tissue organization in breast cancer is both informed by the subtype of the tumour and correlates with patient outcomes^14, 15^.

While two-dimensional, molecularly annotated maps of tissues represent an important advance and clearly provide added value to disaggregated datasets, three-dimensional datasets with layered information comprising as many types of measurements as possible would maximize our ability to extract new biological insights. A recent analysis of gene expression *in situ* during mouse embryogenesis, which includes a time dimension, illustrates the power of such approaches^16^ (*Lohoff et al., Nat Biotechnology (in press)*).

In general, spatial “omics” methods suffer from two main limitations: they usually operate only on thin two-dimensional sections of tissue, and their speed of acquisition is relatively slow. The first issue can be resolved by analysing serial sections cut sequentially from an embedded tissue fragment, and aligning the resulting datasets in order to produce a coherent three-dimensional object. However, the slow acquisition speed means that each individual section can take hours or days to acquire. Multiplied by the large section numbers needed to cover a three-dimensional volume with good axial resolution, this means that a 1×1×1mm block can take many days to weeks in acquisition time alone to produce a dataset.

An additional challenge is the enormous amount of data generated by spatial omics technologies. Spatial omics methods, even when operating in two dimensions, routinely produce datasets containing tens to hundreds of thousands of cells, each possibly described by tens to hundreds of different measurements. Beyond the computational infrastructure necessary to process and store this volume of data, the amount of information produced by these methods can be overwhelming. This issue is heightened for three-dimensional data, since exploration of this data is invariably performed on two-dimensional surfaces. The ability to visualize high-dimensional, complex data is a key element to enabling full exploitation of any spatial molecular profiling strategy. Approaches to address such issues include the use of dimensionality reduction to visualize a dataset in a simplified format or exploitation of machine learning to recognize spatial patterns. However, in both cases, the ability of an individual to explore a dataset naively is greatly diminished. Several software packages exist to facilitate visualization of spatial “omics” datasets, some have arisen as spatial add-ons to widely-used single-cell analysis packages (such as *Seurat*^17^ and *Scanpy*^18^), and some developed from scratch with spatial analysis in mind (i.e. Giotto^19^, Vitescce, ST viewer^20^). However, none of these can handle three-dimensional data natively.

Virtual Reality (VR), and its cousin, Augmented Reality, have significantly matured in the last few years and are set to revolutionize the gaming and entertainment industries. These technologies are increasingly applied in industrial design, medicine, healthcare, and education^21^. However, application of VR to the exploration of scientific data has lagged behind. Indeed, VR has just started to be applied to biological data, mostly to explore gene networks, to annotate neuron morphology, or to visualize microscopy data, both academically^22–25^ (i.e. vLume, CellexalVR, TeraVR, confocalVR) and commercially (Arivis InviewR).

Here, we describe a VR visualization and analysis platform for spatial multi-omics data. This ‘virtual laboratory,’ Theia, so named after the Titaness of sight in Greek mythology, provides a multi-user, three-dimensional environment for exploration of spatial datasets. This platform can accept segmented data with large numbers of molecular or other annotations, project them in 3D, and enable their intuitive manipulation and visualization of patterns of gene or protein expression. Theia is also enabled for on-the-fly analysis, for example with integrated functions for dimensionality reduction that can be used to identify groups of cells for re-projection into the 3D object. Additional analysis capability can be easily added to customize the platform through a convenient Application Programming Interface (API). Theia operates using the Unity VR engine in the Steam VR environment and is therefore compatible with a variety of consumer VR devices and accompanying high-spec personal computers.

To illustrate the utility of Theia for visualization and exploration of 3D datasets, we generated three segmented, molecularly annotated samples (Table 1). Two biopsies from human breast cancers (in the sub-mm size range) were analysed by serial-sectioning followed by IMC and realignment of the processed imaging data. The third dataset is a large-scale model representing a 300 µm thick section of am approximately 1 cm tumour from a syngeneically transplanted mouse triple negative breast cancer (TNBC) model. The latter model was produced by serial blockface two-photon imaging (Serial Two-Photon Tomography, STPT) followed by IMC on recovered sections and contains over 2 million cells.

**Table 1.**
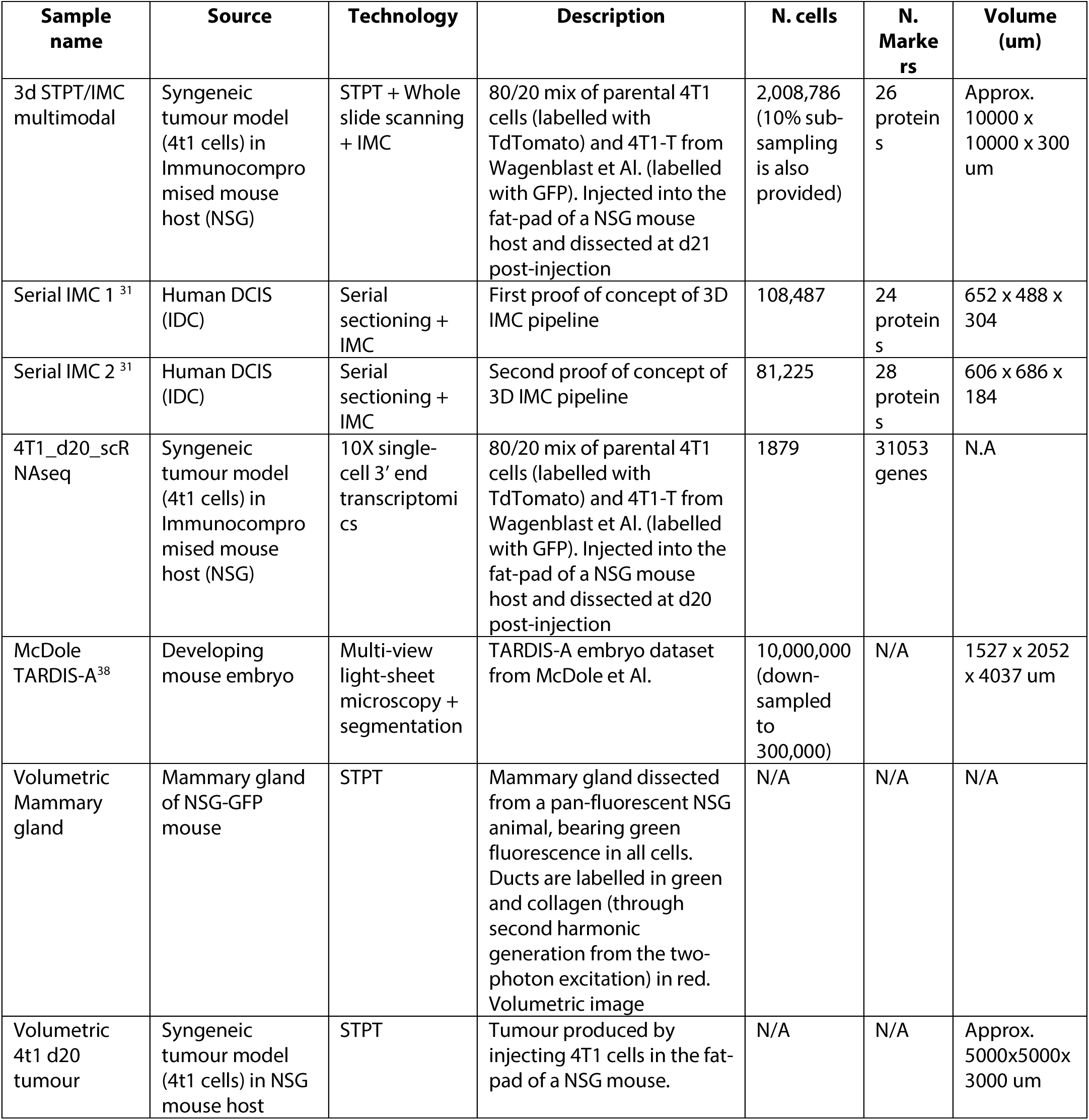
Datasets included for analysis in VR and their source.

While Theia was designed for segmented molecularly annotated datasets, it is also compatible with a wide variety of different data sources, ranging from segmented, 3D lightsheet microscope data (including time-course data) to non-spatially resolved single-cell RNAseq datasets, which can still benefit from the ability to visualize the dimensionality reduction plots in 3D. Any single-cell level data can in principle be visualized in virtual reality, benefitting from the added sense of depth, a simplified user interface, and a large visualization space.

## RESULTS

### The IMAXT pipeline for spatial analysis of biological samples

The *Imaging and Molecular Annotation of Xenografts and Tumours* (IMAXT) CRUK Cancer Grand Challenge project is a collaboration between research groups located in the UK, USA, Canada, Switzerland and Ireland. The goal of this consortium is to enable the production of 3D multi-parametric models of breast tumours in order to increase the current understanding of tumour progression, dissemination, and resistance to therapy, and to pave the way for the development of better diagnostic tools and treatments. At the core of the project is a sample processing and analysis pipeline (*González-Solares, E.A. et al., 2021, Nature Cancer, submitted*) that our consortium developed, combining single-cell genomic/transcriptomic methods with large-volume 3D imaging via STPT and *in situ* molecular profiling (Figure 1).

**Figure 1.**
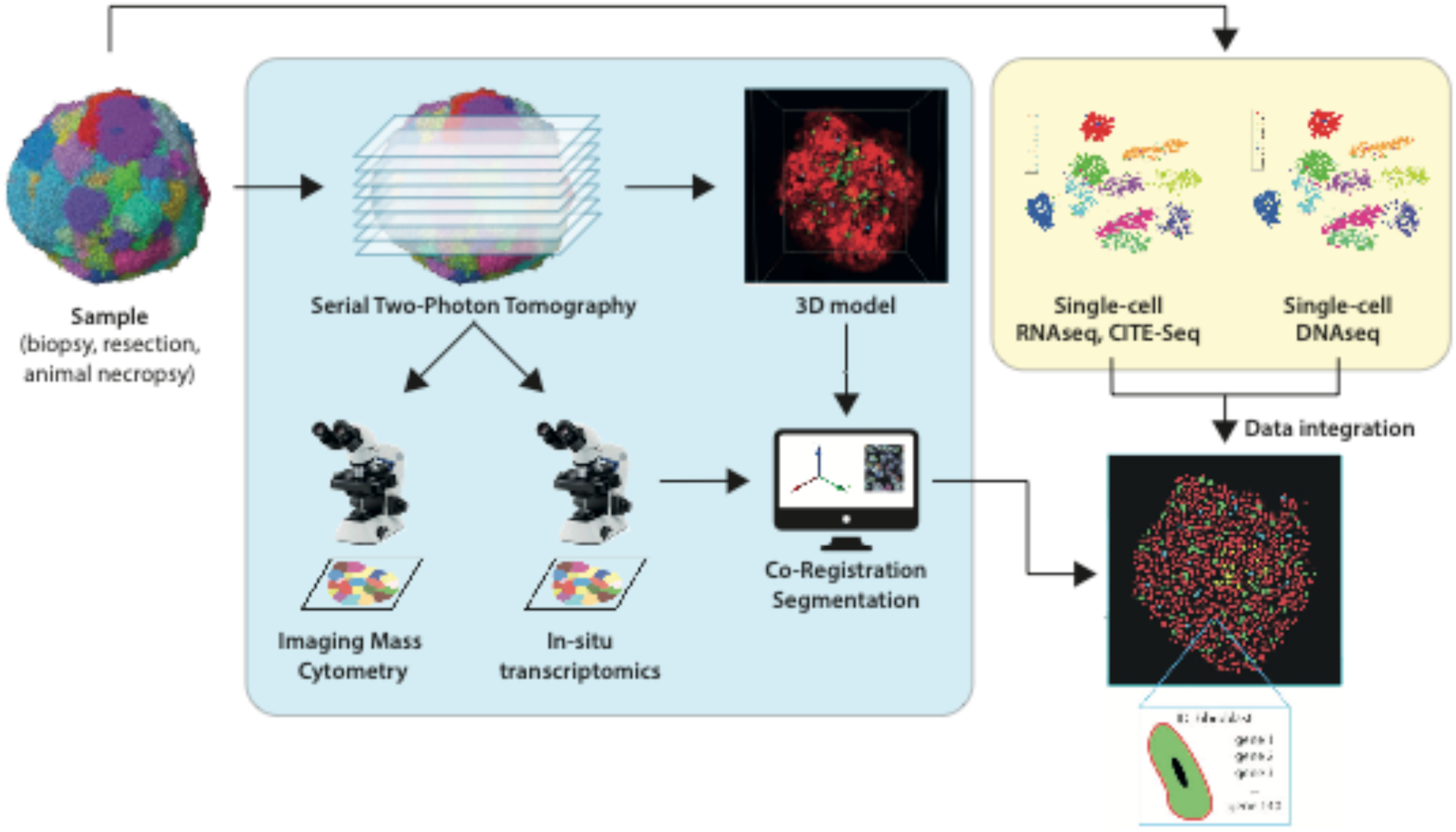
Scheme of the IMAXT pipeline. The IMAXT pipeline integrates data from multiple platforms into a 3D map of a biological sample. Platforms include survey single-cell sequencing, Serial two-photon tomography, Imaging Mass Cytometry, spatial transcriptomics (not included in this manuscript), and data integration.

Samples that enter the pipeline within IMAXT can come from mouse models of breast cancer (obtained from genetically engineered mouse models (GEMMs), from tumour cell lines injected into syngeneic hosts, or chemically or virally induced tumours), patient-derived xenografts grown in immune-compromised mice, or primary human biopsies or resections. As a first step, one or more fragments of each sample are collected and cryo-preserved, and used (after dissociation) for single-cell RNA sequencing (usually using the 10X genomics platform) or single-cell whole-genome DNA sequencing via DLP+ ^26^. The results of these are used to produce a “survey” of the cell types and cell states present in the sample, and to guide the development of the gene and antibody panels used for *in situ* profiling. Since the data yield of disaggregated methods is presently superior to that of *in situ* technologies, this step is very important to ensure the spatial analysis can capture the most relevant parameters, unless a good set of curated markers is already available.

The bulk of the sample is then embedded in an agarose block and subjected to STPT imaging^27^ This method produces a distortion-free 3D image of the sample with up to 4 fluorescence channels. This instrument images a plane immediately below the block surface and then cuts a thin (>=15 μm) section from the sample to enable imaging of the next plane of the sample. In short STPT combines a vibratome and a two-photon microscope. STPT presents three key advantages over other whole-organ images modalities such as light-sheet microscopy: (1) image quality and resolution don’t change with imaging depth, since imaging always happens within a short distance from the top surface of the block; (2) objects of any size and depth can be imaged, with an adjustable axial resolution (physical slice thickness); (3) the sample is sectioned (>=15 μm currently) and imaged at the same time, producing sections that can be further annotated. These gather in a collection vat in the STPT instrument and are recovered (in random order) on glass slides or in a known order with an automated collection device (currently limited to sections of >25 μm). In our workflow, sections are counter-stained with nuclear and membrane dyes and re-imaged using a whole slide scanner. Each slide image is then registered to the 3D volume produced by two-photon imaging. To facilitate this step, we co-embed spherical agarose beads coated with green fluorescent protein in the sample block. The cross-section of the beads produces a unique pattern for each slice, which can be easily segmented and matched across imaging modalities (*González-Solares, E.A. et al., 2021, Nature Cancer, submitted*). Collected sections are then available for further molecular annotation using spatial transcriptomic or proteomic methods.

### Generation of an integrated dataset from a mouse model of TNBC using STPT and IMC

The 4T1 cell line is a widely used model for basal-like, claudin low breast cancer^28, 29^. It can be syngeneically transplanted in immune-competent Balb/C animals or introduced into immune-compromised recipient mice. Prior work has indicated that 4T1 is both genomically and phenotypically heterogeneous, and we previously identified subclonal populations of 4T1 (clones E and T) which can selectively form vessel-like networks when orthotopically transplanted^30^. We therefore used this cell line to create a demonstration model for Theia.

We fluorescently labelled the parental 4T1 cell line by expression of TdTomato and labelled the 4T1-T subclone by expression of GFP. We orthotopically injected a mixture containing 80% parental and 20% clone T into an NSG recipient animal and resected the tumour after 21 days, when its diameter had reached approximately 1 cm. We imaged the entire tumour using the STPT and collected 20 consecutive sections for further annotation, corresponding to a depth of approximately 300 μm. These sections were imaged on a slide scanner, stained with a panel of 33 metal-conjugated antibodies and imaged by IMC. After co-registration and segmentation, we obtained a 3D model with an axial resolution of 15 μm (roughly an individual cell diameter) and over 2,000,000 unique cells mapped over a volume of approximately 12 mm^3^.

Even in the 3D STPT image alone, we noted different spatial distribution for clone T as compared to the parental 4T1 cells (Figure 2A, Supplementary video 1). The former distributed in “foci” within the tumour mass, while the latter were largely uniform. Being able to combine this imaging with IMC revealed that clone T gathered often in proximity to necrotic areas with substantial myeloid infiltration (Figure 2 B-D, S1, Supplementary video 2). Interestingly, tumour cells expressing the proliferation marker Ki67 often had low expression of the TdTomato fluorophore (Figure 2F). We could also identify host vascular cells expressing endothelial markers (CD31/CD34) and pericyte markers (PDGFRB, Desmin, alpha-SMA), stromal cells, and cell clusters characterized by high expression of the M2 macrophage marker CD206/MMR (Figure 2 C,E, Figure S1). In addition to cell types, we were also able to interrogate cell states. We could clearly identify hypoxic cells, both in the TdTomato+ tumour compartment and in the GFP+ one, as well as cells characterized by high amounts of phosphorylated S6 (Ser235/236) ribosomal protein, a downstream target of the mTOR pathway (Figure 2F).

**Figure 2.**
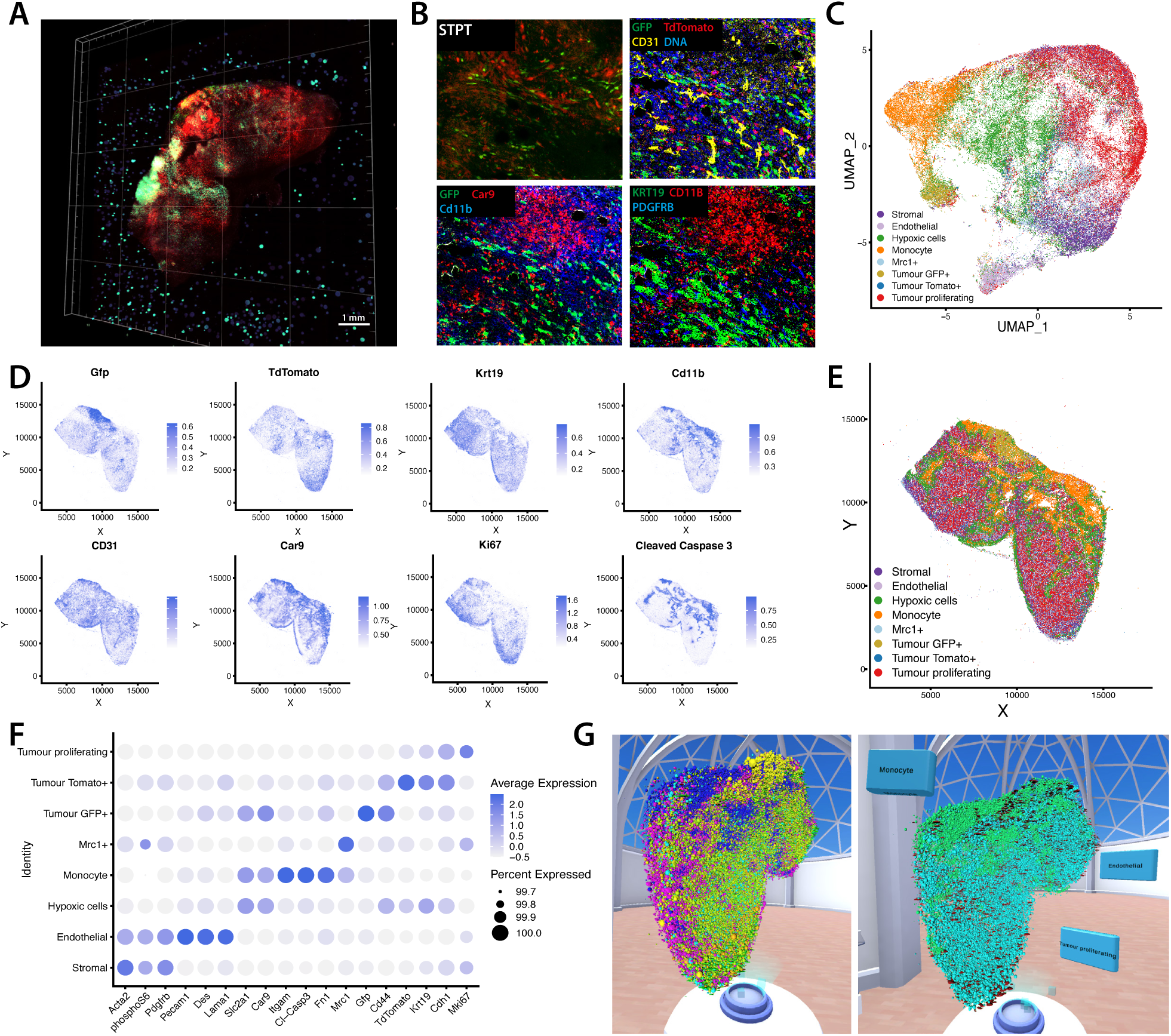
Mutimodal analysis in 3D of a 4T1-derived tumour. **A.** 3D view of the STPT data for the multi-modal NSG 4T1 model dataset. GFP is displayed in green and TdTomato in red. **B.** Zoom-in on an area of the sample including a vessel-like structure. Cutouts display the full resolution STPT data, as well as IMC images for GFP/TdTomato (tumour cell populations), immune markers (Cd11b), Hypoxia markers (Car9), Vessel markers (CD31), Epithelial markers (Krt19), Pericyte markers (PDGFRB), and nuclear counter-stain. **C.** UMAP dimensional reduction plot for the dataset. **D.** Marker abundance plots for a representative 2D section of the sample. Each dot is a segmented cells and signal intensity corresponds to the normalized abundance of the marker in the cell. The same markers described above are displayed, with the addition of proliferation (Ki67) and apoptosis (Cleaved Caspase 3). **E.** Spatial plot of a representative 2D section of the dataset. Each colour corresponds to a different cell type predicted by leiden clustering on the data. **F.** Dot plot identifying specific markers for each cell population. Signal intensity corresponds to average expression, dot size corresponds to the fraction of the population expressing the marker. **G.** 3D model as visualized in Theia. All cells are shown on the left (Endothelial cells: red - Hypoxic cells: pink – Myeloid cells: blue - Mrc1+ cells: cyan – Stromal cells: green - Tumour GFP+ cells: yellow - Tumour Tomato+ cells: orange, Tumour Ki67+ proliferating cells: lime), and specific cell types are shown on the right (myeloid cells in green, proliferating tumour cells in cyan and endothelial cells in red).

Overall, our results indicated a clear spatial organization of the tumour, with distinct populations of tumour cells occupying discrete neighbourhoods. In general, GFP+ neighbourhoods were characterized by low proliferation, low expression of epithelial markers, and high levels of hypoxia-related proteins (I.e. Glut1, Carbonic Anhydrase IX). These cells also had higher expression of CD44 than the parental 4T1 cells (Figure 2F). All of these characteristics can be interrogated in 3D in virtual reality using Theia (Figure 2G, see below).

### High-resolution 3D datasets from human cancer by serial sectioning and IMC

As additional input to our virtual reality platform, we have also provided two human breast cancer samples that have been previously generated as part of IMAXT Consortium efforts. These models were generated from a Her2+ invasive ductal breast carcinoma biopsy using 3D IMC to achieve high axial-resolution together with *in situ* molecular annotation (*Kuett and Catena et al. Nature Cancer - submitted*)^31^. Briefly, the 3D IMC method is based on serially sectioning a paraffin-embedded tissue sample into 2 µm sections prior to sample staining and acquisition. This is followed by computational image processing to achieve the final 3D model with a voxel size of 1×1×2 µm (x-y-z). The first model has the final dimensions of 652 x 488 x 304 µm^3^, and the second one 606 x 686 x 184 µm^3^ and comprise 108,000 and 81,000 segmented cells, respectively (Figure 3A, left and right respectively), providing an interesting substrate for exploration in VR (Figure 3B-F). Notably, these models reveal spatial distributions of marker expression that are not evident in 2D images, thus illustrating the ability of 3D models to raise interesting hypotheses. For example, these models show a varying pattern of CK5-high and SMA-high epithelial basal cells, a dense colocalization of CD8+ and CD4+ T-cells near endothelial cells, and lining of pS6(Ser235/236)-high cells along the epithelial tumour compartments (Figure 3C-E, S3, Supplementary video 3). In the larger 3D IMC model, clusters of potentially invasive cells at the tumour periphery have been captured, and the 3D visualization makes it apparent that these clusters are disconnected from the main tumour bulk (Figure 3F).

**Figure 3.**
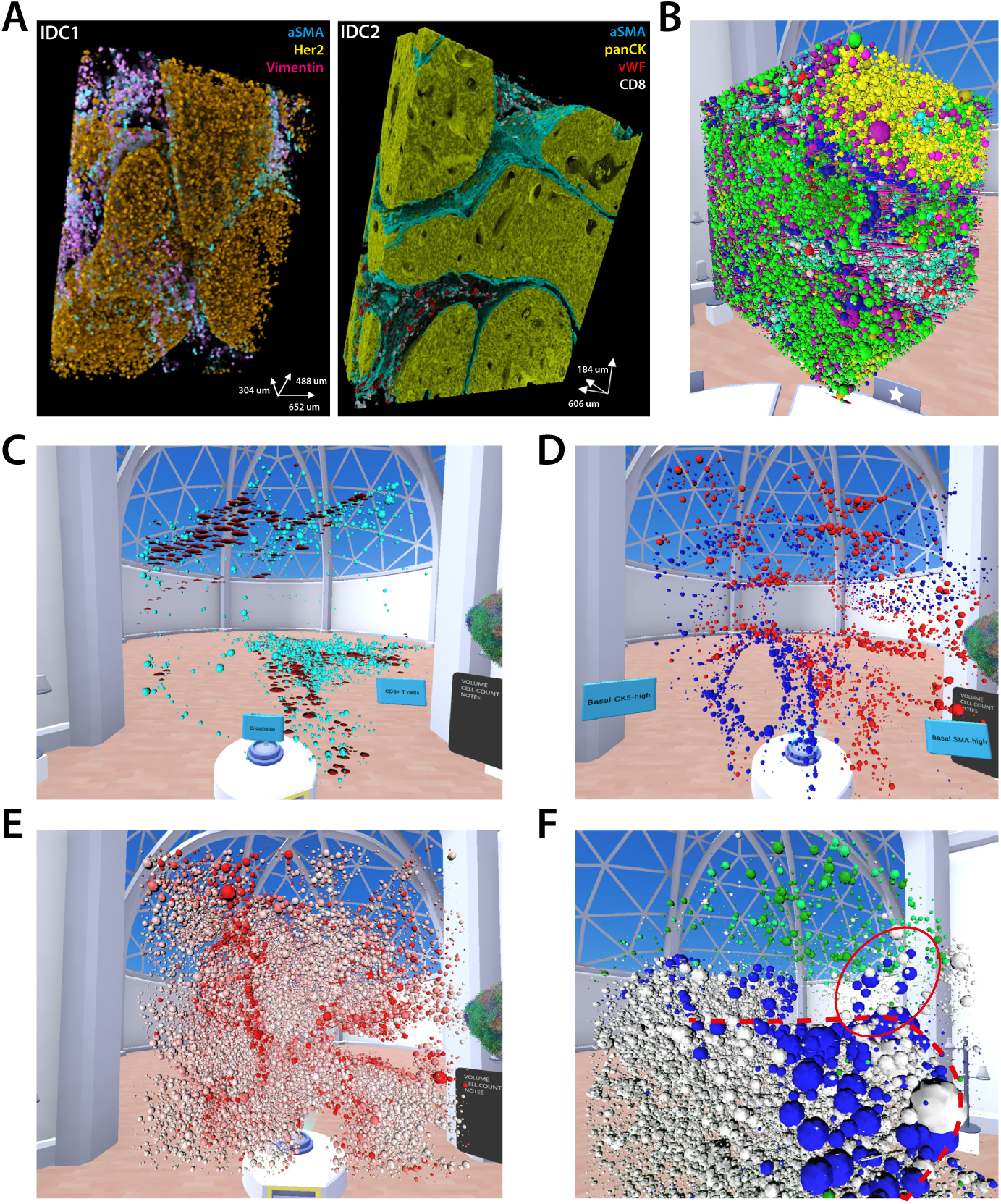
VR analysis of Invasive ductal carcinoma models generated by serial sectioning IMC by Kuett, Catena et al. **A.** 3D renderings of the raw IMC data for the first and second model, showing (as indicated) basal markers (SMA), Stromal markers (Vimentin), epithelial markers (panCK / Her2), endothelial markers (vWF/CD31) and T cell markers (CD8). **B.** IDC2 model visualized in the VR environment in Theia (the Z scale is exaggerated to facilitate visualization). **C.** CD8+ T cells (cyan) clustered around endothelial cells lining a blood vessel (red) visualized in the VR model. **D.** Lining of tumour-filled mammary ducts formed by CK5+ and SMA+ basal cells (in red and blue). E: Intensity of phospho-S6 (Ser235/236) marker visualized through virtual reality, highlighting a pattern of increased expression just below the outer surface (basal layer) of the ducts. **F.** IDC1 visualized in VR. Some tumour cells from a specific phenograph cluster (in blue, red circle) can be seen breaking off a mammary duct (red dotted line) and invading the stroma (in this case rich in lymphocyte cells – green). Other tumour cells are in white.

### Development of a virtual reality ‘laboratory’ to explore spatial molecular data

To enable exploration of three-dimensional, molecularly annotated datasets, we developed a virtual reality implementation that takes lessons from the computer gaming industry and applies these with the goal of enabling intuitive spatial analyses of cellular arrangements and gene expression patterns in biological samples. VR has the advantage of not being bound by the dimensions of even a very large computer monitor. Instead, users can literally step inside the data, generally in a roughly 5m^2^ area and apply different analysis paradigms and explore the outcomes on the fly. Finally, our implementation creates a multi-user environment, such that investigators can convene in a virtual space to interrogate collaboratively high dimensional datasets. We named this implementation, Theia. Theia has the potential to impact the analysis of spatial data analogously to the impact of the development of genome browsers have had on our ability to interrogate and understand next-generation sequencing datasets.

Theia was developed on consumer-grade VR headsets (any compatible with the SteamVR system) and computer hardware to make it accessible to any laboratory. The investment needed for operating the software, including headset, controllers and computer is less than the cost of a generating one single-cell dataset. The software is built upon a popular and well-supported game engine, Unity, which results in broad compatibility. Our primary concern was to create a space that was comfortable to work in for several hours with tools that responded quickly and usefully. Additionally, it was imperative for us that no programming be required and data analysis be achieved with simple visual metaphors, to make the learning curve as shallow as possible for new users. The objective was a balanced design, both scientifically functional and with limited visual clutter (Figure 4A, Supplementary Videos 4-5).

**Figure 4.**
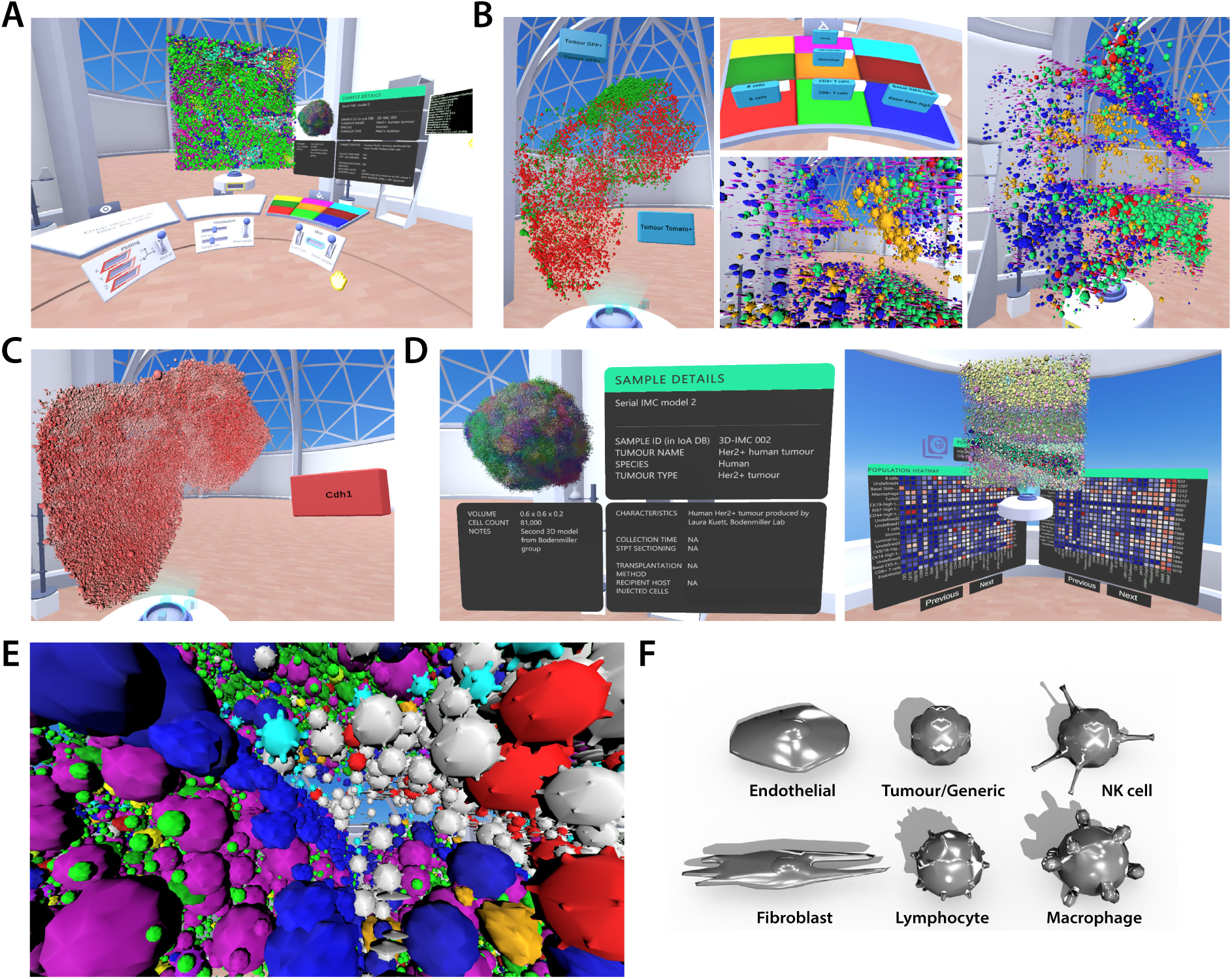
Data exploration tools in Theia. **A.** Overview of the virtual environment featuring the selection table and plotting slots (left), favourites table and sample/cell size console (centre), and sample information panel, highlighting table and export console (right). **B.** Cell type selection for the NSG 4T1 model (GFP+ and TdTomato+ tumour cells displayed) and of the IDC2 3D-IMC model (detail of the highlighting table and 3D visualization with stroma in magenta, macrophages in orange, B cells in red, CD8+ T cells in green, and Basal SMA+ cells in blue. **C.** Marker intensity mapping for E-cadherin on the NSG 4T1 model. **D.** Detail of the sample information panel (left) and population information panel and expression heatmap (right). **E.** A fly-in of the IDC2 model. The camera is located in a stromal region just outside a duct (blue cells, bottom left) filled with tumour cells. Other cell types are represented in different colours (i.e. green/pink tumour cells, red immune cells) **F.** Meshes representing different cell types in icon view.

Theia tools have a universal language which is easy to learn. They consist of two varieties; surfaces for generic analysis (displaying and highlighting cell populations, or visualizing gene expression) and cabinets for specific analysis such as dimensional reduction or differential expression. Grabbing and placing of objects is the main mode of interaction. For instance, the sample can be rotated and zoomed just by grabbing it and moving it closer or farther away Supplementary videos 4-5). Additionally, pointing, buttons, sliders, and levers are utilized to change visualization parameters such as cell size and scale. Tools for moving, selecting specific cells, pointing and taking screenshots are housed on the user’s wrists (Figure S2A). This ensures easy access from anywhere within the simulation. Features of the dataset to be analysed (populations, genes, cell types) are displayed as VR ‘tiles’ which the user can interact with by moving them in specific slots into the environment tools (See examples in Figures 4, 5, Supplementary videos 6-7. These interactions are subtly guided by objects “snapping” together when in proximity of a compatible slot, and by haptic feedback through the VR controllers. Subtle animations and in world screens are also utilized within Theia. This prompts the user toward correct actions and ensures simulations become intuitive (Figure S2B, Supplementary video 8).

**Figure 5.**
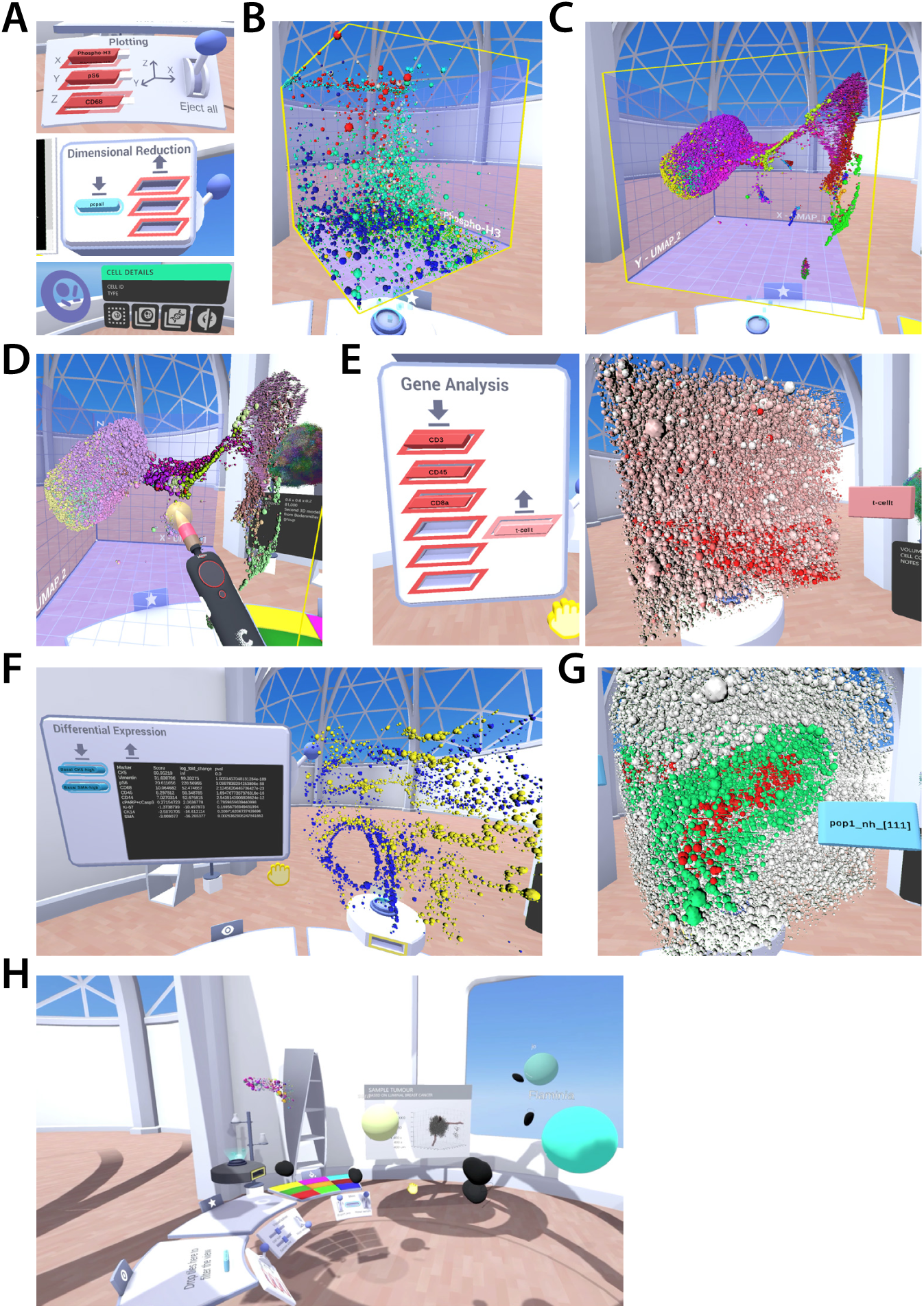
Analysis tools in Theia. **A.** detail of the gene plotting tool, dimensional reduction tool, and population manipulation tool. **B.** example 3D marker expression view for phospho-S6, (Ser235/236) Phospho-Histone H3 and CD68. Different cell clusters are highlighted in different colours (i.e. macrophages in red) **C.** 3D UMAP plot of the IDC2 sample. **D.** Cell selection tool. Selected cells are highlighted in the centre of the model. **E.** Gene signature tool and intensity mapping of a T-cell signature produced by averaging the per-cell intensities of CD3, CD8a, and CD45. **F.** Differential expression tool showing DE genes for the CK5^hi^ or SMA^hi^ populations of basal cells (in yellow and blue). P values are generated by T-test. **G.** Neighbourhood search tool. An initial selected population (in red) is expanded by a user selected radius to generate the green area. **H.** Multi-user interaction with Theia: 3 users analysing a sample together. Note the head avatars and the controller objects tracking the users’ hands.

### Interactive data analysis in the VR environment

Theia is designed to analyse samples formatted as cell catalogues, with each cell characterized by a series of spatial coordinates (corresponding to either real space or embeddings along dimensional reduction axes) and parameters (gene/protein expression or morphological measurements). Cells can be pre-assigned to different populations (for instance by means of clustering methods such as the louvain or leiden algorithms^32, 33^), or populations can be defined directly by the user by “painting” over cells with a specific tool. Theia uses a dedicated data storage format optimized for small size and speed of loading, but can import data from most single-cell processing data formats, e.g., the one specified by the *anndata* Python package used by *Scanpy*^18^, or R’s *singlecellexperiment* and *Seurat*^17^.

Cell populations can be made to appear or disappear by selecting the tiles corresponding to these cell types from a menu and dropping them on a selection table and can be assigned a colour by dropping them on a second table (Figure 4B, Supplementary videos 6-7). All cell populations can be automatically displayed in sequence (Supplementary videos 9-10). Genes and markers can be visualized in a similar way. Dropping a gene tile on the selection table launches a query to identify cells with an expression of the given gene above or below a give threshold (which can be selected) (Supplementary video 11), while dropping it onto a colour re-maps the colour intensity to the gene expression level, with a transfer function that can be personalised adjusting the min/max expression range (Figure 4C, Supplementary video 12). Queries can be combined by dropping multiple tiles on the selection table, and multiple markers or populations can be assigned colours at the same time.

Information about the sample is displayed through dedicated “panels” which display details at the sample level (general metadata on sample generation, size, features, etc) (Figure 4D), or at the population level, or single cell level. For the latter two, the panel shows a customizable maker expression heatmap, which allows users to get a general view of gene or protein expression in a population or in a given cell. The information displayed in the panels changes dynamically as the user selects different cells (Figure 4D, Supplementary video 13).

While most of the manipulations happen with the model mapped to a size (in virtual space) more or less equal to that of the user (so that it can be held and manipulated), it is also possible for the user to greatly expand the model and use a “fly” tool, colloquially termed “superman mode,” to enter the sample and interrogate individual cells and structures (Figure 4E, Supplementary Videos 14-15).

Although data on the 3D shape of each cell is available in our datasets, we elected to display each cell using simplified meshes corresponding to the most frequently observed cell types (icon view) (Figure 4F). This ensures the user can visually parse the sample with ease and also provides an additional layer of visual information. Furthermore, icon view is computationally simpler to render, and allows for the viewing of larger samples. The icon assigned to each different cell type can be defined when samples are imported. In spatial datasets such as the ones produced by our group, in which information on the volume of each cell is present, the mesh size of each object is taken from the real measurements. In other data types, the size can be used as a visual cue for a parameter, for instance by making specific cell types appear bigger or smaller to facilitate their visualization.

We implemented tools to switch easily between visualization of the spatial coordinates of the sample and the expression of specific genes or embeddings produced by dimensional reduction algorithms. Gene/marker tiles can be inserted in a “plotter” tool, which transforms the sample visualization into a 3D plot with dimensions corresponding to the marker intensity (Figure 5B, Supplementary video 16). The same can be done with dimensional reduction embeddings, which can either be pre-generated in the dataset or calculated on the fly (UMAP reduction^34^ is used by default). (Figure 5C, supplementary video 17) Users can perform a full “round trip” analysis, for instance selecting a spatial area for study, visualizing the cells contained in it in gene space or in a dimensional reduction plot, identifying a population and mapping it back to its spatial location (Figure 5D, Supplementary video 18). Other tools allow the definition of “gene signatures” (by averaging the expression of multiple markers into a new marker tile that can be used as a single parameter) (Figure 5E) or the calculation of differential gene/marker expression between custom populations (Figure 5F, Supplementary video 19). Finally, a spatial search tool can be used to identify all cells lying within a certain distance (in 3D) from a selected population (Figure 5G, Supplementary Video 20). This function is critical to identify patterns of marker expression that depend on spatial proximity between cells (i.e. by comparing differential expression between cells closer or farther away from a specific tissue feature). Cell populations identified from within Theia can be exported as csv files which can be further processed through any other processing pipeline.

All of the functions of Theia are available in multi-user environments hosted on public servers. Users can host or join interactive analysis sessions in which the same dataset is analysed cooperatively, and up to 8 international users have been convened in a single session thus far. Each user is represented by an avatar in the analysis space, and can interact with the objects in the virtual laboratory. Audio feeds enable live communication between users during analysis (Figure 5H, Supplementary video 21).

### Advanced analysis and plug-in structure

In designing Theia, we aimed to strike a balance between implementing easy-to-use tools directly in the software, and leveraging the incredibly rich ecosystem of single-cell analysis software, scripts, and methods (some of which are dedicated to spatial analysis) that already exist and are in constant evolution. For this reason, one of the key features of Theia is a plug-in system which allows users to pass information between the visualization software and a custom piece of code, run some processing, and return data for visualization. The viewer component of Theia (developed in Unity and C# language) sits on top of a Python instance. The dataset loaded in the viewer is automatically also loaded in Python as an *anndata* object, and is available for manipulation using any Python module, such as the *scanpy* package. Users can develop custom scripts performing their desired processing, and, by using specific tools in the virtual space, load it and input parameters (such as list of cells) to it. The script output is then parsed and returned to Theia in the form of new cell populations, new markers, or gene lists, and can be displayed in VR. This method is used, for instance, for the dimensionality reduction, differential expression, and spatial search tool, which can be customised or varied by the expert user by modifying the underlying Python code. New scripts can also be developed from scratch to implement other functions. Different templates (corresponding to different cabinet shapes in the virtual world) mediate different types of input-output, and can be used to implement different functions. While at the moment the plug-in engine is only compatible with Python, we plan to extend it to R in the near future, and it is already possible to call R scripts by using Python as an intermediate.

### Compatibility of Theia with other data types

We expect spatial datasets, such as the ones presented here, to become increasingly available. However, true, molecularly annotated, 3D datasets are still relatively rare and difficult to generate, requiring substantial infrastructure and effort in technology implementation and data analysis. We therefore have also made Theia compatible with the analysis of datasets and datatypes that are already widespread.

Disaggregated single cell analyses (transcriptome, copy number, methylome, etc) are increasingly used in all areas of biology, including to interrogate the tumour microenvironment and the clonal evolution of different tumour types. The technology to generate many of these data types is now mature and broadly available. Theia is compatible with all single-cell datasets, and includes a data converter for scanpy’s *anndata* objects, one of the most popular formats. Other formats can be loaded if converted first to *anndata*, which is usually a straightforward process. If positional information is not included, the dimensional reduction embeddings (t-SNE, PCA or UMAP) are used in lieu of the spatial coordinates for visualization, and populations/clusters are mapped to cell types. All of Theia’s tools (on-the-fly dimensional reduction, differential expression, gene averages, etc) are compatible with single-cell data. As an example of the usage of this data type, we provide with Theia a single-cell dataset obtained from the same tumour type and stage represented in our STPT/IMC 3D model (Figure 6, Supplementary video 22)

**Figure 6.**
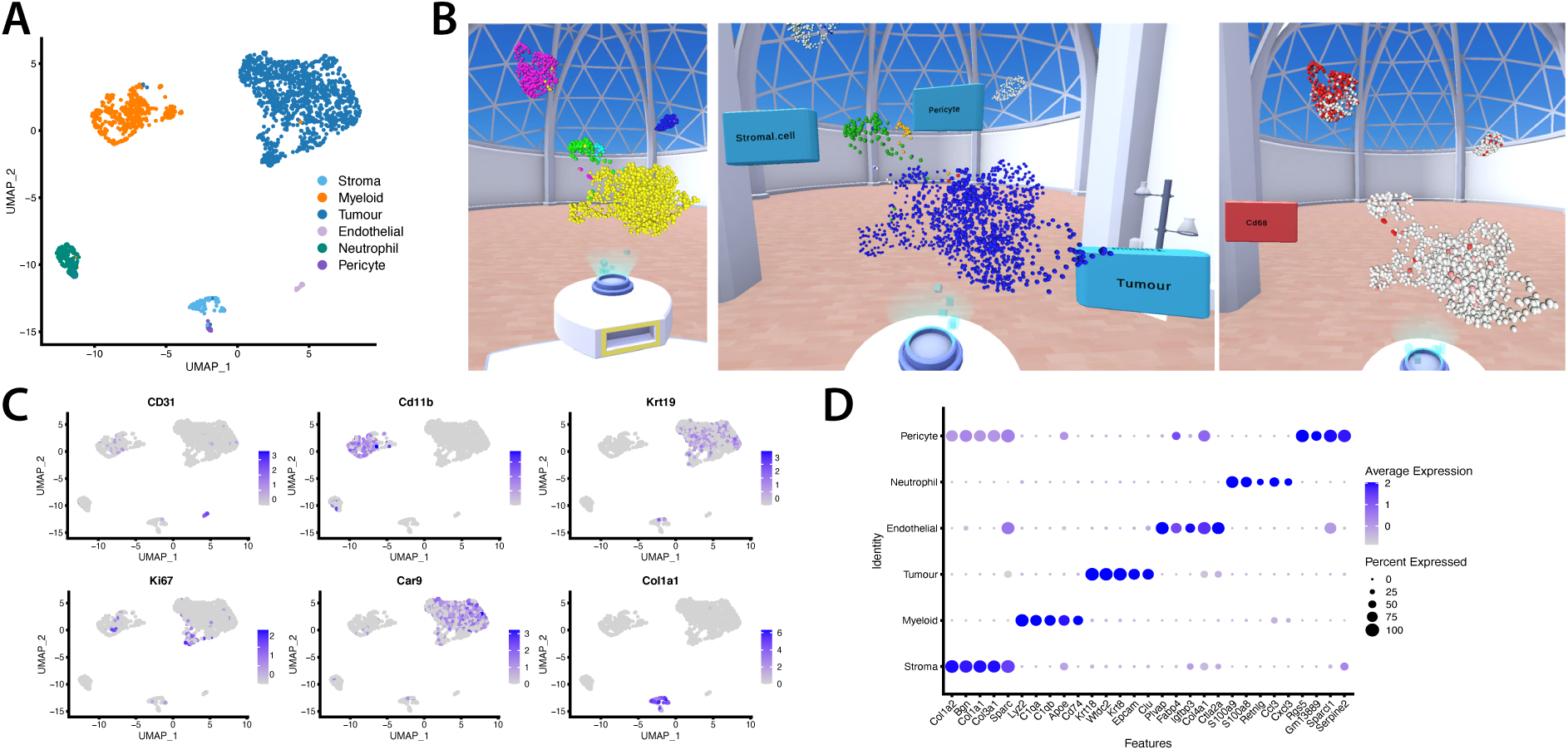
Visualization of disaggregated single-cell datasets in Theia. The dataset displayed here was generated through the 10X genomic platform (3’ GEX v3) from a NSG 4T1 tumour analogous to the one used for the STPT/IMC dataset. Approximately 1000 cells are included **A.** UMAP plot highlighting cell types as inferred by the *cellassign* supervised cell identification algorithm. **B.** Model shown in Theia. All cells (left), cell highlighting (centre) and gene intensity mapping (right). **C.** UMAP plots showing marker expression for a subset of markers also present in the STPT/IMC dataset and for the stromal marker Col1a1 (collagen 1). **D.** Dot plot identifying specific genes for each cell population. Signal intensity corresponds to average expression, dot size corresponds to the fraction of the population expressing the marker

Whole-mount microscopy is another family of technologies that has recently seen increased democratization, in part thanks to the development of novel microscopy techniques such as light sheet, bessel beam, lattice light sheet, STPT, and others^35^. Thanks to technical advancements, whole organisms can now be imaged *in toto*, including live imaging of their development. Datasets of this type have been published for *D. melanogaster* and zebrafish early development^36, 37^ and for pre-implantation mouse embryos during gastrulation and early organogenesis^38^. In parallel to the technical improvements in imaging, improved methods for object segmentation have allowed the identification of every cell in these models, and their tracking across time. Since the output of these processes is a catalogue XYZ position for each cell at different timepoints, Theia can be used to visualize them. To enable visualization of time-series data, we modified one of the functions of our viewer, normally used to cycle through the different cell types present in a sample to be compatible with timepoints. As a result, Theia can be used to load and visualize spatial time-lapse data. Cell labels and markers (if present) can still be visualized as for other samples. As an example of this, we included with Theia a pre-packaged version of the mouse development dataset from McDole et al. (Figure 7A, supplementary Video 23) for exploration.

**Figure 7.**
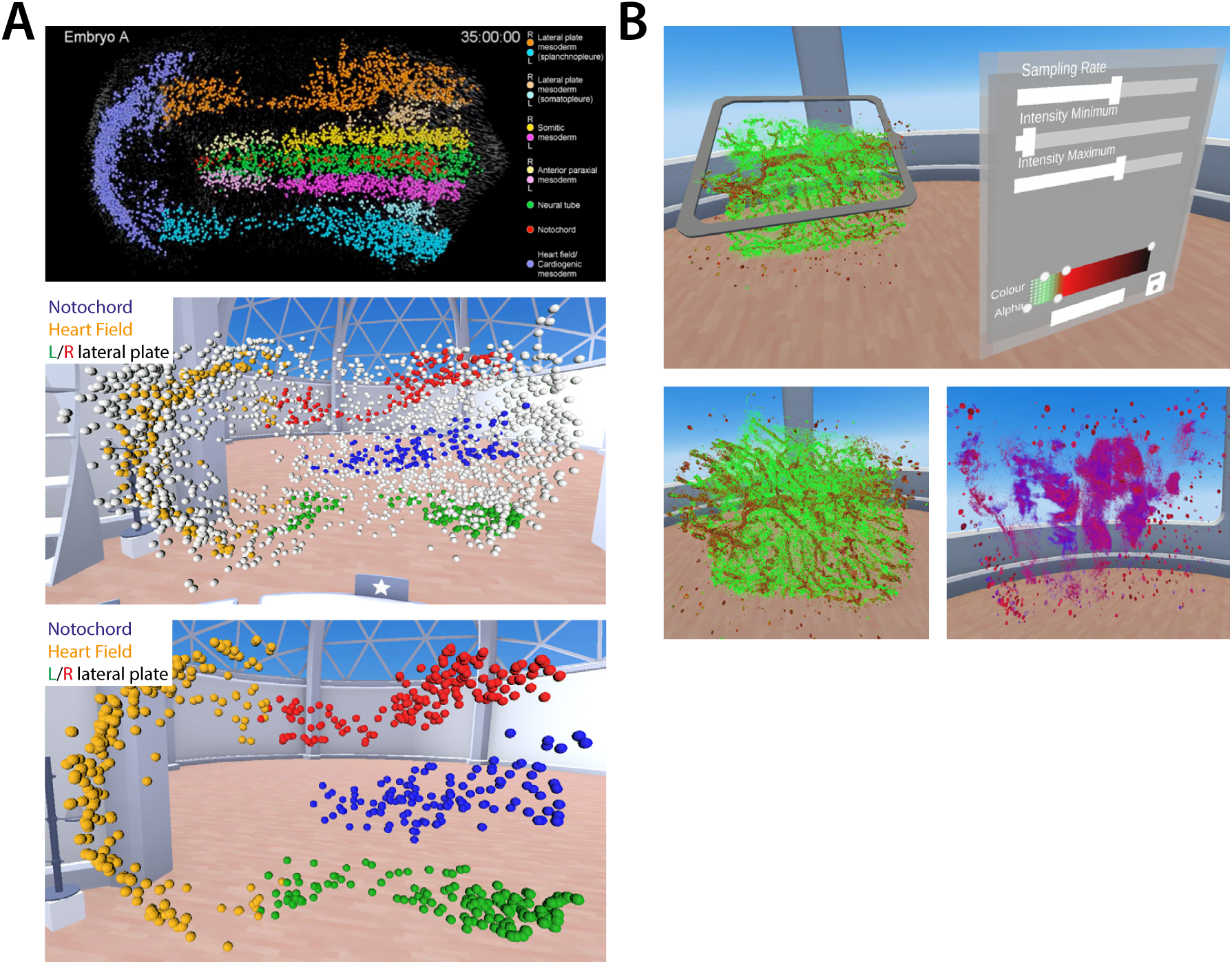
Compatibility of Theia with other data types. **A.** The embryo neurulation dataset was from McDole et al ^38^ Top: modified figure from McDole et Al showing cell localizations and lineages 35h after the start of imaging. Middle/bottom: Theia visualizations of the same dataset. The neural tube, lateral plates and heart fields are highlighted. **B.** Theia volumetric viewer for the visualization of native voxel data. Top and bottom-left: Example mammary gland dataset generated by STPT on a fragment of a GFP+ mouse mammary gland. All mouse tissues are GFP+. Collagen detected via second harmonic generation is in red. The middle panel showcases the sectional plane tool. Bottom right: 3D model of a 4T1 tumour.

Finally, while segmentation of biological images and production of annotated cell catalogues is arguably a critical step of most analysis pipelines, we sought to make Theia compatible with non-segmented data as well, and introduced a volumetric renderer into the software. While this is not yet capable of visualizing terabyte-scale datasets at native resolution (mostly due to limitations in data I/O and to the fact that Unity’s renderer is not natively designed to handle voxel data), it is sufficient to visualize image volumes up to many millimetres in size with a resolution in the 10s of microns, sufficient to discriminate minute features of the tissue microenvironment. The volume viewer takes as input tiff image stacks, and provides controls to adjust the transfer function, opacity, colour and sampling density of the visualization. The viewer is by default compatible with single-colour images, but will be soon expanded to multi-channel images. Users are able to move and zoom volume around by grabbing it and manipulating it, and are provided with a “plane” tool which allows them to “section” the volume at any level and angle and display that sectional plane. The latter operation, which is normally quite laborious on a computer’s screen, and requires long manipulations of the dataset to get the correct angle and position, can be performed in seconds in virtual reality, which further highlights the power of this medium for volume analysis (Figure 7B, Supplementary video 24).

## DISCUSSION

The increasing availability of commercial platforms for producing spatial omics data is poised to impact biology in a manner similar to the widespread deployment of commercial single-cell profiling methodologies. This necessitates the development of platforms to analyse, process, and explore such datasets. Virtual Reality offers a flexible, scalable, and intuitive solution for the exploration of spatial data, and this has motivated the creation of Theia as an open-source resource for the community. Theia has been created with embedded tutorials (accessible from within the VR environment) to enable investigators to rapidly master its use, and most users can gain a familiarity with Theia’s basic toolkit in under 30 minutes. Theia also offers a suite of more advanced analyses and importantly the ability to integrate most existing tools and analyses for single-cell level data and provides the ability to easily incorporate novel tools in the future. Moreover, Theia has been developed to be compatible with widely available and relatively inexpensive hardware, with a complete installation generally costing less than producing one single-cell dataset.

## METHODS

### Sample generation for the 3D/STPT sample

To construct the tdTomato-Akaluc vector, tdTomato and p2A-Akaluc were amplified by PCR and cloned into the 3rd generation lentiviral pZIP backbone harbouring a spleen focus-forming virus 57 promoter (SFFV) using Gibson Assembly. 4T1 parental cells (ATCC® CRL-2539™) were infected with tdTomato-Akaluc followed by selection of tdTomato positive cells using FACS. 4T1-T cells (as described in Wagenblast et al.^30^) were made GFP positive by transduction with the CellTag-GFP vector (pSMAL60 CellTag-V1), which was a gift from Samantha Morris^39^ (Addgene 115643), based on the pSMAL backbone from John Dick, and were sorted based on their EGFP expression. The cell lines were cultured in DMEM (Gibco) supplemented with 10 % (v/v) heat-inactivated FBS in a humidified incubator at 37°C and 5 % CO_2_.The murine tumour 4T1 parental cells (tdTomato^+^) were combined with the 4T1-T clonal cells (EGFP^+^) in a ratio of 80 % parental-4T1 to 20 % 4T1-T. From this mixed population, 50,000 cells were suspended in 1:1 PBS and Matrigel (Corning, Cat No. 356231) and tumour formation was induced by orthotopic injection into the mammary fat pad of female NSG™ mice (Jackson labs, NOD.Cg-Prkdcscid Il2rgtm1Wjl/SzJ, stock 005557). Primary tumour volume was assessed and the tumour was allowed to develop for 21 days post-injection before the animal was sacrificed and the tumour was excised. The tumour was fixed in 4 % paraformaldehyde (PFA) for 24 h, 4°C. All of the animal procedures were performed in accordance with the UK Animal Scientific Procedures Act (ASPA) under the authority of an animal use project license approved by the UK home office, and in accordance with the standard operating procedures indicated by CRUK CI Animal Welfare Review Board (AWERB).

### Fiducial beads conjugation for STPT embedding

NHS-activated Sepharose® beads (Sigma Cat No. GE-17-0906-01), composed of 4 % cross-linked agarose with an average particle size of 90 µm, were coated with recombinant GFP protein (Abcam, Cat No. Ab84191). Briefly, an aliquot of 200 µL bead slurry was washed in 1 mM HCl, followed by Coupling Buffer (0.4 M NaHCO_3_, 1 M NaCl, pH 8.3). 100 µg of GFP was incubated with the beads overnight at 4°C, with rotation. The reaction was quenched with 0.1 M Tris-HCl pH 8.0, 0.3 M NaCl and the GFP-conjugated beads were stored in 0.1 M Tris, pH 8.0 at 4°C.

### STPT sample preparation and STPT imaging

To prepare the tumour sample for serial two photon tomography (STPT), a 4.5 % (w/v) agarose solution (Type 1 agarose, Cat. No. A6013, Sigma) in 50 mM phosphate buffer, pH 7.4 was oxidized by the addition of 10 mM sodium periodate (NaIO_4_). The solution was agitated for 3 h in the dark, washed with phosphate buffer and resuspended in the appropriate volume of 50 mM phosphate buffer to achieve 4.5 % agarose. This oxidized agarose solution was heated to boiling, cooled to 60°C, and spiked with GFP-conjugated agarose fiducial marker beads prior to embedding the tissue. The PFA-fixed tissue was embedded in the molten agarose using a 2 cm^3^ embedding mould (Cat. No. E6032-1CS, Sigma). Once the agarose block was solidified, it was immersed into a degassed polymer solution (Imbed 100S monomer, TissueVision) for 48 h at 4°C. The agarose block was baked for 8 h at 40°C to crosslink the polymer solution, following which the block was stored at 4°C in 50 mM phosphate buffer, pH 7.4 until it was ready for imaging.

The agarose block containing the tumour sample was glued to a histology glass slide (25 x 75 x 1 mm) modified by attaching two neodymium bar magnets to the bottom side (non-frosted) using epoxy glue, and this was placed onto a magnetic plate within an imaging vat and filled with 1 lt of 50 mM phosphate buffer, pH 7.4. Serial two-photon imaging was then performed on a TissueCyte 1000 instrument (TissueVision, Newton, MA, USA), where a series of 2D XY mosaic images were taken, followed by physical sectioning with a vibratome to remove the imaged tissue and to create a new surface for a subsequent round of imaging. For this dataset, twenty physical sections of 15 µm thickness were cut with the vibratome at a speed of 0.1 mm/sec and 55 Hz frequency. Two planes of images were taken for each 15 µm physical section, one at an imaging depth of 30 µm below the surface and another at 38 µm. A dual laser setup (Coherent Discovery) allowed simultaneous acquisition of GFP (excitation wavelength 900 nm) and tdTomato (excitation wavelength 1040 nm). Fluorescence was detected by four PMT tubes in the following spectral ranges: <500nm (channel 4), 500-<u>560</u> nm (channel 3), 560-600nm (channel 2), >600nm (channel 1). GFP and TdTomato were detected in channel 3 and channel 2 respectively. Collagen was also imaged through the second-harmonic emission generated by the 900nm laser (450 nm). Tissue sections were collected from the buffer vat onto Superfrost Plus microscope slides, air-dried and stored at 4°C until processed for fluorescence scanning. The tiled STPT images were stitched and segmented using the image analysis workflow described below, and in more detail elsewhere (*González-Solares, E.A. et al., 2021, Nature Cancer, submitted*).

### Slide imaging

Images were captured post-STPT with the Zeiss Axioscan Z1 microscope slide scanner with a resolution of 0.44 µm/pixel. GFP - Excitation filter – 465-490nm, Emission filter – 460-480nm, Exposure time 250ms. tdTomato – Excitation filter – 545-565nm, Emission filter – 578-640nm, Exposure time 700ms. The surface area of each section was imaged including the GFP+ fiducial marker beads surrounding the tissue.

### IMC antibody conjugation and panel preparation

All antibody conjugations were performed using the standard protocol available from Fluidigm for metal-antibody conjugation using the Maxpar X8 metal conjugation kit. All centrifugations were done at room temperature. Briefly, the metal polymer was equilibrated to room temperature and spun down in a mini-centrifuge for 10 seconds. The polymer was suspended in 95 µl of Fluidigm’s L buffer and 5 µl of the appropriate lanthanide metal from Fluidigm was added to this. After thorough resuspension, the metal-polymer was incubated at 37°C for 30 minutes. During this step, 100 µg of an IgG antibody (in BSA and glycerol free formulation) were spun down at 12,000 x g for 10 minutes in an Amicon 50 kDa centrifuge filter tube. A buffer exchange was performed by adding 400 µl of Fluidigm’s R buffer to the concentrated antibody and spun down at 12,000 x g for 10 minutes. The antibody was then partially reduced with 100ul of 4mM TCEP-R buffer, made by diluting 0.5 mM TCEP (Sigma-Aldrich) in R-buffer. After ensuring gentle and thorough resuspension of the antibody and reducing agent, the antibody was incubated in a water bath at 37°C for 30 minutes. During this step, the polymer-lanthanide mixture was transferred to an Amicon 3 kDa centrifuge filter tube with 200 ul of L buffer and spun down at 12,000 x g for 25 minutes. The polymer-metal complex was then washed with 400 µl of Fluidigm’s C buffer and spun down at 12,000 x g for 30 minutes. During the polymer-metal purification, 300 ul of C buffer was added to the partially reduced antibody and spun down at 12,000 x g for 10 minutes. This wash was repeated with 400 µl of C buffer. After purification of both the polymer-metal complex and the reduced antibody, the polymer-metal was brought to a final volume of 80 µl with C buffer and added to the 20 µl of antibody, thus initiating the conjugation processes. The antibody-polymer-metal complex was incubated in a water bath at 37°C for 90minutes. The conjugated product was then washed with 200 µl of Fluidigm’s W buffer and spun down at 12,000 x g for 10 minutes. The wash was repeated 3 times more with W buffer up to a total volume of 400 µl. The ∼20 µl of the conjugated antibody was resuspended in W-buffer to a final volume of 100 µl and its absorbance at 289 nm was measured using the Nanodrop. After spinning down the product at 12,000 x g for 10 minutes, the conjugated antibody was resuspended in the appropriate amount of PBS to bring the final concentration to 0.5 mg/ml. The product was spun down at 1,000 x g for 2minutes, collected and supplemented with 0.05% sodium azide preservative. Final conjugated products were stored at 4°C for long term use. The final panel used for this manuscript is described in Supplementary Table 1.

### IMC sample preparation for the STPT/IMC dataset

IMC imaging was performed on 20 consecutive 15 µm STPT sections adhered to frosted glass slides. Briefly, sections were incubated at 60°C for 30 minutes, followed by a 5-minute wash in ddH_2_O. Slides were placed into 50-ml Falcon tubes containing antigen retrieval reagent (Tris-EDTA pH9), preheated at 95°C in a water bath. Slides were transferred to the pre-heated ARR for 30 minutes, and subsequently cooled under running cold water for 5 minutes to ensure gradual reduction to 70°C. Slides were then washed in ddH_2_O for 10 minutes followed by TBS (tris-buffered saline) for 10 minutes. Tissue sections were first permeabilised in a 0.3% Triton X-100/TBS buffer for one hour, then blocked in a 3%BSA/0.3%Triton X-100/TBS solution for one hour. The blocking solution was removed, ensuring removal of excess liquid to avoid diluting the antibody mix. The antibody mix (Supplementary Table 1) was prepared in a final solution of 1%BSA/TBS and added to each tissue section; a coverslip was placed onto each tissue section. The slides were placed in a humidified chamber at 4°C for overnight incubation. The following day, the coverslips were gently removed and slides were washed twice in 0.1%Tween 20/TBS, followed by two washes with TBS, each wash performed for 7 minutes. The tissue section was then stained with Fluidigm’s DNA intercalator (catalogue #201192B), dilute 1:500 in TBS, and incubated for 30 minutes at room temperature. Sections were washed with ddH_2_O for 5 minutes and allowed to air dry before imaging.

### IMC Image acquisition

Each tissue section was imaged using the Hyperion Imaging System™ (Fluidigm). The system was first tuned and calibrated using a glass slide that’s been labelled with known concentrations of each metal isotope within Fluidigm’s metal library. Calibration was performed at a frequency of 20 Hz and an ablation energy of 0 dB, with Pre-Calibration XY Optimization and Fine XY Optimization options both enabled. Upon successful calibration of the system, the slide was placed into the ablation chamber and a panorama image of the entire tissue section, including the surrounding STPT, GFP-labelled beads, was generated. Acquisition was performed at a frequency of 200 Hz and an ablation energy of ranging from 0 to 5 dB depending on acquisition time and laser duty hours (energy was calibrated to the lowest amount sufficient to produce complete ablation on a small tissue sample). Tuning and image acquisition ablations were all performed using a UV laser set a diameter of 1 µm. The data acquired was stored into both an MCD and txt file.

### STPT image stitching and Z alignment

Because STPT is the modality that has access to a relatively intact sample cube, STPT images serve as the anchor for data processing. It was important then to ensure that these images provide the most accurate representation of the sample possible.

With our choice of focal lenses, the field of view of the STPT microscope is roughly 1 mm^2^, with a pixel size of 0.56 μm. Imaging the surface of a sample cube requires typically around a hundred field of view acquisitions; we refer to these as tiles. The control software for the microscope has the capability to reconstruct the full stage view from these tiles by stitching them together from the recorded positions of the actuators that move the sample. We have found that these positions can be often be 10 μm off, and therefore chose to implement our own stitching procedure (*González-Solares, E.A. et al., 2021, Nature Cancer, submitted)*.

For this purpose, we configured the STPT microscope so that there is a 10% overlap between adjacent tiles, and we refined the tile-to-tile positioning by intensity-matching these overlaps. This allowed us to reconstruct the full stage image with relative tile positioning errors of around 1 μm.

The normal operation of the microscope consists of an acquisition in several channels and/or optical depths, followed by sectioning the sample with the microtome. After this, the sample cube is raised, and the cycle begins again. Because the sample is re-positioned between slices and each slice image is stitched independently, there are mis-alignments introduced between slices. These could be corrected by means again of comparing slice to slice, but because the sample itself has changed, this is often imprecise. To solve this problem and facilitate multi-modal registration later on, we used the embedded Sepharose® beads as fiducial marks.

These beads are spherical and therefore easy to segment and model. For the former, we used a U-Net neural network^40^ trained on manually segmented beads. For the latter, we built a realistic model of a homogeneous sphere embedded in a medium of transmissivity T<1.0. From this model we derived the centre coordinates and the radius of each bead. This effectively transformed the beads into point sources, allowing us to resort to a wealth of algorithms developed for Astronomy, as aligning/crossmatching point sources is a common problem in this field.

In the case of slice-to-slice registration, a simple nearest neighbour search was enough to find the bead pairs in contiguous slices, and by identifying in which slices a given bead appeared (due to their size, usually 3 to 5 slices) we could use the measurements of their central coordinates to align all the slices in the sample. Typical alignment errors were of 3 µm, while the action of the microtome introduces a drift that from top to bottom would skew the misaligned cube close to 20 μm in one direction.

### STPT to whole slide scanning to IMC registration

The collection of slices from the STPT tank effectively randomises their order, but once deposited on a glass slide, an image was taken using a Zeiss Axioscan and the slide was given an unique ID that enables traceability. The issue that remained was to find the match between each Axioscan image and the corresponding STPT slice. This was complicated by several factors: firstly, depositing the slice onto the glass slide can change the left/right, top/bottom and front/back orientation; secondly, STPT images a layer embedded some microns into the tissue, while Axioscan does so to the surface of the sample; thirdly, collecting and depositing the slices introduces some mechanical deformation. Our matching algorithm attempted to address all of these factors. First, we aligned a 32x down-sampled Axioscan image to a STPT dummy image obtained from a median along the Z axis of all the slices. We modelled the STPT to Axioscan transformation as an affine transform with six degrees of freedom, solving the problem of relative orientation and correcting most of the possible deformations. We then segmented and profiled the visible beads on the Axioscan image following the same procedure as for STPT, and using the rough transformation derived previously we tried to match the detected Axioscan beads to those in each STPT slice. Maximising the number of beads in common gave us the best Axioscan to STPT match, along with the STPT to Axioscan transformation for that particular slice.

Once a sample slice was affixed to a glass slide and tagged, the IMC to Axioscan correspondence was immediate. As IMC imaging is time consuming, smaller stage sizes are used, and normally fewer beads are visible. Having Axioscan as a middle stage alleviates the associated problems. We registered each IMC multichannel cube to its associated Axioscan image using the same procedure outlined before, saving the first step as in this case relative orientation is fixed. Once a good transformation was obtained, we compounded it with the already known Axioscan to STPT transform, and refined this by comparing the beads in common between Axioscan and STPT.

Typical registration errors, as measured by comparing the bead centers are of 6 μm for the IMC to STPT registration and 7 μm for the Axio to STPT one.

### IMC segmentation

IMC images were segmented using an automated pipeline developed for high-throughput analysis of biological images. The pipeline is written in Python and uses the OpenCV library, i.e., an open-source computer vision and machine learning software library written in C++. For each IMC slice, the pipeline reads a data cube, performs a pre-processing step, segments individual cells, extracts several features for each cell, and finally outputs a catalogue of detected cells and their calculated properties as well as a cell mask image. Initially, the code extracts the nuclear channel using metadata information and uses this channel as a reference for the subsequent segmentation process. It includes the normalisation of the reference image and noise reduction by applying a Gaussian filter. Next, we used an adaptive threshold method to remove background pixels. The latter produced a binary image which was the input to the watershed segmentation algorithm. Most of the cells overlapped with one another. Therefore, one crucial step was to separate (also called deblending) such cells. Our algorithm used the coordinates of local peaks (maxima) to perform the deblending task. At this step, we computed cellular features for each segmented cell. These included centroids, shape descriptors, and mean pixel intensities within the cell nuclei across all available IMC channels using the cell nuclear mask. In addition, we created another image mask associated with the cell’s periphery, i.e. the two-dimensional zone surrounding the cell, to compute mean pixel intensities (across all IMC channels) within the cytoplasmic area. Finally, we produced a catalogue that includes all properties for detected cells and a cell image mask for all detections. The pipeline is also further described in a separate manuscript (*González-Solares, E.A. et al., 2021, Nature Cancer, submitted)*

### Dataset assembly and single cell analysis for the STPT/IMC model

In order to produce a coherent 3D dataset, the cell catalogues produced by segmentation of each IMC dataset were first transformed into the coordinate frame of the STPT 3D model by applying the transformation defined above (STPT to Axio to IMC registration) to the XY coordinates of each cell, and using the progressive order of the STPT section matched to the IMC image as the Z. This produced a three-dimensional set of cell localizations, which was transformed into an *Anndata* python object for further processing using the *scanpy* package. In order to clean the dataset, any IMC channel corresponding to antibodies that did not produce a good quality staining pattern, or that weren’t specific for a pathway or cell type (i.e. the nuclear counterstain) were removed.

An initial dimensional reduction (using the *pca*, *neighbour* and *umap* functions of the scanpy package) was performed for the initial IMC section, followed by clustering using the *leiden* function. The cells from the subsequent sections were then aligned (in multi-dimensional parameter space) with the annotated ones from the first section using the *ingest* function, re-projecting them into the same UMAP coordinate space and transferring the cluster labels. Clusters were then manually annotated and cleaned based on user experience and the expression of known cell type markers. Finally, a new 3-dimensional UMAP reduction was calculated for the entire dataset. The final dataset was saved in *Anndata* format as well as converted to *Seurat* format for wider compatibility.

### 3D IMC human breast cancer models

The two 3D IMC human breast cancer models were used after final pre-processing as presented in Kuett and Catena et al. (2021, Nature Cancer, submitted). The data was downloaded from https://doi.org/10.5281/zenodo.4752030. Briefly, for the 3D IMC models, paraffin embedded biopsy samples were serially sectioned into 2um sections using diamond knife and ultramicrotome. After section collection samples were stained with a mixture of metal-tagged antibodies and acquired with a commercial Fluidigm Hyperion Imaging System. The raw mcd image files were converted into omeTIFF files using IMC pre-processing pipeline available at https://github.com/BodenmillerGroup/imctools. Consecutive images were aligned using Fiji-ImageJ2-linux64 v1.0 plugins Register Virtual Stack Slices and Transform Virtual Stack Slices (https://github.com/fiji/register_virtual_stack_slices/). 3D segmentation was done with a Fiji-ImageJ2-linux64 v1.0 plugin called h-watershed (https://github.com/mpicbg-scicomp/Interactive-H-Watershed/).

### Single cell dataset processing

The single cell sample distributed with Theia was produced from a tumour similar to the one described above in “Sample generation for the 3D-STPT sample”. In short, an 80/20 mixture of parental 4T1 cells (not fluorescent) and green-labelled 4T1-T cells (Zsgreen) were injected into the fat pad of two Nod-Scid-Gamma (NSG) mice (Jackson labs, NOD.Cg-Prkdcscid Il2rgtm1Wjl/SzJ, stock 005557). Tumours were allowed to develop for 20 days and collected by necropsy. Tumour were dissociated using a Miltenyi Biotech GentleMACS octo w/heaters dissociatior and the Tumour dissociation kit (cat. 130096730) according to supplier’s instructions. Cells were washed, counted, and approximately 4000 cells per sample were submitted to the CRUK-CI genomics core for processing through the 10X genomics 3’ single cell gene expression pipeline (v3). The sequencing results were processed using the 10X *Cellranger* software to produce cell-gene count matrices, and the resulting datasets were further processed using the *Scanpy* package.

Approximately 1800 cells were identified as high-quality after removing doublets, cells with low counts, cells with high mitochondrial reads and cells with high ribosomal reads. Celltypes were identified using the *cellassign* R package^41^ and a custom annotated marker matrix. Each cluster was further validated by performing unbiases clustering with the *leiden* method and verifying that each cellassign cluster corresponded to one or more leiden clusters. Normalization, log-transformation, scaling and dimensionality reduction/neighbourhood analysis were all performed using the *scanpy* package.

### Mammary duct volumetric dataset

The mammary gland sample was obtained from a virgin NSG-GFP mouse (Jackson Labs, *NOD.Cg-Prkdcscid Il2rgtm1Wjl Tg(CAG-EGFP)1Osb/SzJ*, stock 021937, approximately 2-3 months old. The animal was first euthanised using increasing concentration of CO2 as prescribed by the schedule 1 of the UK Animal Scientific Procedures Act (ASPA) and specific in the home office approved project license mentioned above. The inguinal fat pad was dissected and fixed for 24h in 4% PFA in PBS, followed by several washed in PBS. The sample was embedded and imaged as described in “STPT sample preparation and STPT imaging” above, with the following variations: 8% agarose was used, and the sample was infiltrated with Imbed 301H+ monomer mix instead of Imbed 100S. STPT imaging was performed taking 100 15um sections, GFP was visualized in channel 3 and second harmonic generation (collagen) in channel 4. Excitation was 900nm from the tuneable beam of the Coherent Discovery laser attached to the STPT instrument.

### Packaging of samples for VR analysis

Datasets in *Anndata* format can be converted directly into the proprietary format used by Theia by means of a python script provided with the Theia software (anndata_to_theia.py). The script guides the user through a series of questions aimed at defining the coordinate sets to use as spatial coordinate (i.e. spatial coordinates or dimensional reductions), the cell type annotation to use, the variables to associate to size, which icons should be associated to different cell types, whether to normalize data, etc.

For the IDC samples downloaded from Zenodo and for the embryo development sample from McDole et Al, the conversion was performed by means of custom Jupyter notebooks available on the github site mentioned below.

### Theia software

Theia is built with the Unity real time 3D development platform. The choice was made to build upon an existing 3D framework over creating one from scratch. This would allow for a shorter development time as well as the access to support when issues arose. Unity in particular was chosen for its flexible nature as well as the abundance of developers and artists that are familiar with it. Unity’s multiplatform nature allowed us to minimise risk, as it builds on top of existing standard modules (the SteamVR platform) supporting multiple devices. It is also a very fast tool for prototyping and experimentation, which would be an important aspect for developing such a new piece of software. Theia is primarily programmed in C#. Python is used to import, analyse and export datasets.

In order to display so many 3D objects in virtual reality we needed to use instanced rendering. We used this to allow us to display large volumes of the same mesh. This allows us to show larger samples in the simulation. In addition to this we use compute shaders for highlighting of the sample. Using a technique called parallel reduction we are able to maintain a smooth framerate and retain comfort in the simulation while making each cell an interactable object. Framerate is critical in virtual reality, as low values lead to the user experiencing motion sickness. For this reason, we employed several strategies to minimize rendering load during fast interaction, for instance applying a “vignette” effect blanking the periphery of the field of view during fly movements.

A Python socket server was used to connect with external Python tools and execute custom scripts on the sample. This was used for differential expression, dimensionality reduction and the nearest neighbour function. These were implemented using the *scanpy* package.

### Data and code availability

IMAXT aims to produce periodic data releases for the scientific community. The data described in this manuscript have been included in our Imaxt Data Release 1 (IDR1). This release contains the raw STPT, whole-slide scanning and IMC data for all models, as well as the re-projected whole-slide and IMC data, the segmentation masks, the cell catalogues, and the final dataset in *anndata* and *seeurat* format. Information on how to access the data and description of datasets can be found in https://imaxt.ast.cam.ac.uk/release/docs/dr1/). IDR1 also contains the single-cell RNAseq dataset described in figure 6, which is also deposited in the Gene Expression Omnibus (GEO), with ID GSE178069. The data for the serial ablation IMC human IDC models can be obtained from the Zenodo platform https://doi.org/10.5281/zenodo.4752030. The code used to perform stitching of the STPT sample and nuclear segmentation of the IMC datasets is released under a GNU General Public License version 3 (GPLv3) and publicly available from https://github.com/IMAXT/ (*González-Solares, E.A. et al., 2021, Nature Cancer, submitted)*. Executables and source code for Theia can be downloaded from https://imaxt.ast.cam.ac.uk/release/docs/dr1/ and from https://www.suil.ie/

## Supporting information

List of consortium authors

## ACKNOWLEDGEMENTS

We wish to acknowledge the core facilities of the CRUK-Cambridge Institute. In particular we thank Light Microscopy (Stefanie Reichelt), Flow Cytometry (Richard Grenfell), Genomics (Paul Coupland, Kania Katarzyna), Histopathology (Cara Brodie, Julia Jones), and Research Instrumentation and Cell Services (Jane Gray). Furthermore, we wish to acknowledge the CRUK-CI laboratory management team, IT team, and Biological Resources Unit (BRU) for their hard work and assistance during this project. We wish to acknowledge Clare A. Rebbeck for her work in managing the ASPA (UK Animal Scientific Procedures Act) Project License under which all the animal work described in this manuscript was performed, and for her suggestions and feedback while designing animal experiments. The IMAXT VR Tiger Team (Owen Harris, Robert Becker, Sara Vogl, Flaminia Grimaldi, Suvi Coffey, Dario Bressan, Mohammad Al Sa’d, Giorgia Battistoni, Ilaria Falciatori, Dan Goodwin, Laura Kuett, Ignacio Vazquez-Garcia, Joanna Waselak, Spencer Watson, Jonas Windhager) provided critical feedback during the development project as did the wider IMAXT collaboration. The fluorescently labelled cell lines used to produce the 3D STPT/IMC model were generated by Sophia Wild (IMAXT laboratory). The CellTag-GFP vector (pSMAL60 CellTag-V1) was a kind gift from Samantha Morris (Addgene 115643). This work was supported by Cancer Research UK [A24042, A21143]. G.J.H. is a Royal Society Wolfson Research Professor.

## AUTHOR CONTRIBUTION

*Project design and overall direction:* Dario Bressan, Owen Harris. Gregory J. Hannon

*Data generation:* Dario Bressan, Claire M. Mulvey, Fatime Qosaj, Laura Kuett, Raul Catena, Ilaria Falciatori

*Data analysis:* Dario Bressan Ali Dariush, Carlos Gonzalez-Fernandez. Eduardo A. Gonzalez-Solares, Mohammad Al SA’d, Aybuke Kupku Yoldas, Tristan Whitmarsh

*VR software development*: Robby Becker, Flaminia Grimaldi, Suvi Coffey, Sara Lisa Vogl, Owen Harris

*Manuscript preparation:* Dario Bressan, Fatime Qosaj, Claire M.Mulvey, Eduardo A. Gonzalez-Solares, Carlos Gonzalez-Fernandez, Ali Dariush, Nicolas Walton, Bernd Bodenmiller, Owen Harris, Gregory J. Hannon

## LITERATURE CITED

1. Joyce, J. A. & Fearon, D. T. T cell exclusion, immune privilege, and the tumor microenvironment. Science 348, 74–80 (2015).

2. Quail, D. F. & Joyce, J. A. Microenvironmental regulation of tumor progression and metastasis. Nat. Med. 19, 1423–1437 (2013).

3. Yuan, Y. Spatial Heterogeneity in the Tumor Microenvironment. Cold Spring Harb. Perspect. Med. 6, (2016).

4. Chen, K. H., Boettiger, A. N., Moffitt, J. R., Wang, S. & Zhuang, X. RNA imaging. Spatially resolved, highly multiplexed RNA profiling in single cells. Science 348, aaa6090 (2015).

5. Moffitt, J. R. et al. Molecular, spatial, and functional single-cell profiling of the hypothalamic preoptic region. Science 362, (2018).

6. Wang, X. et al. Three-dimensional intact-tissue sequencing of single-cell transcriptional states. Science 361, eaat5691 (2018).

7. Lubeck, E., Coskun, A. F., Zhiyentayev, T., Ahmad, M. & Cai, L. Single-cell *in situ* RNA profiling by sequential hybridization. Nat. Methods 11, 360–361 (2014).

8. Ståhl, P. L. et al. Visualization and analysis of gene expression in tissue sections by spatial transcriptomics. Science 353, 78–82 (2016).

9. Rodriques, S. G. et al. Slide-seq: A scalable technology for measuring genome-wide expression at high spatial resolution. Science 363, 1463–1467 (2019).

10. Giesen, C. et al. Highly multiplexed imaging of tumor tissues with subcellular resolution by mass cytometry. Nat. Methods 11, 417–422 (2014).

11. Goltsev, Y. et al. Deep Profiling of Mouse Splenic Architecture with CODEX Multiplexed Imaging. Cell 174, 968–981.e15 (2018).

12. Lin, J.-R., Fallahi-Sichani, M., Chen, J.-Y. & Sorger, P. K. Cyclic Immunofluorescence (CycIF), A Highly Multiplexed Method for Single-cell Imaging. Curr. Protoc. Chem. Biol. 8, 251–264 (2016).

13. Gut, G., Herrmann, M. D. & Pelkmans, L. Multiplexed protein maps link subcellular organization to cellular states. Science 361, eaar7042 (2018).

14. Ali, H. R. et al. Imaging mass cytometry and multiplatform genomics define the phenogenomic landscape of breast cancer. *Nat*. Cancer 1, 163–175 (2020).

15. Georgopoulou, D. et al. Landscapes of cellular phenotypic diversity in breast cancer xenografts and their impact on drug response. Nat. Commun. 12, 1998 (2021).

16. Pijuan-Sala, B. et al. A single-cell molecular map of mouse gastrulation and early organogenesis. Nature 566, 490–495 (2019).

17. Stuart, T. et al. Comprehensive Integration of Single-Cell Data. Cell 177, 1888–1902.e21 (2019).

18. Wolf, F. A., Angerer, P. & Theis, F. J. SCANPY: large-scale single-cell gene expression data analysis. Genome Biol. 19, 15 (2018).

19. Dries, R. et al. Giotto: a toolbox for integrative analysis and visualization of spatial expression data. Genome Biol. 22, 78 (2021).

20. Fernández Navarro, J., Lundeberg, J. & Ståhl, P. L. ST viewer: a tool for analysis and visualization of spatial transcriptomics datasets. Bioinformatics 35, 1058–1060 (2019).

21. Implementing Virtual and Augmented Reality Tools for Radiology Education and Training, Communication, and Clinical Care | Radiology. https://pubs.rsna.org/doi/10.1148/radiol.2019182210.

22. Legetth, O. et al. CellexalVR: A virtual reality platform to visualise and analyse single-cell data. bioRxiv 329102 (2020) doi:10.1101/329102.

23. Spark, A. et al. v LUME: 3D virtual reality for single-molecule localization microscopy. Nat. Methods 17, 1097–1099 (2020).

24. Wang, Y. et al. TeraVR empowers precise reconstruction of complete 3-D neuronal morphology in the whole brain. Nat. Commun. 10, 3474 (2019).

25. Stefani, C., Lacy-Hulbert, A. & Skillman, T. ConfocalVR: Immersive Visualization for Confocal Microscopy. J. Mol. Biol. 430, 4028–4035 (2018).

26. Laks, E. et al. Clonal Decomposition and DNA Replication States Defined by Scaled Single-Cell Genome Sequencing. Cell 179, 1207–1221.e22 (2019).

27. Ragan, T. et al. Serial two-photon tomography for automated ex vivo mouse brain imaging. Nat. Methods 9, 255–258 (2012).

28. Dexter, D. L. et al. Heterogeneity of tumor cells from a single mouse mammary tumor. Cancer Res. 38, 3174–3181 (1978).

29. Aslakson, C. J. & Miller, F. R. Selective events in the metastatic process defined by analysis of the sequential dissemination of subpopulations of a mouse mammary tumor. Cancer Res. 52, 1399–1405 (1992).

30. Wagenblast, E. et al. A model of breast cancer heterogeneity reveals vascular mimicry as a driver of metastasis. Nature 520, 358–362 (2015).

31. Catena, R. et al. Highly multiplexed molecular and cellular mapping of breast cancer tissue in three dimensions using mass tomography. bioRxiv 2020.05.24.113571 (2020) doi:10.1101/2020.05.24.113571.

32. Traag, V. A., Waltman, L. & van Eck, N. J. From Louvain to Leiden: guaranteeing well-connected communities. Sci. Rep. 9, 5233 (2019).

33. Blondel, V. D., Guillaume, J.-L., Lambiotte, R. & Lefebvre, E. Fast unfolding of communities in large networks. (2008) doi:10.1088/1742-5468/2008/10/P10008.

34. McInnes, L., Healy, J. & Melville, J. UMAP: Uniform Manifold Approximation and Projection for Dimension Reduction. ArXiv180203426 Cs Stat (2018).

35. Susaki, E. A. & Ueda, H. R. Whole-body and Whole-Organ Clearing and Imaging Techniques with Single-Cell Resolution: Toward Organism-Level Systems Biology in Mammals. Cell Chem. Biol. 23, 137–157 (2016).

36. Huisken, J., Swoger, J., Del Bene, F., Wittbrodt, J. & Stelzer, E. H. K. Optical sectioning deep inside live embryos by selective plane illumination microscopy. Science 305, 1007– 1009 (2004).

37. Keller, P. J., Schmidt, A. D., Wittbrodt, J. & Stelzer, E. H. K. Reconstruction of Zebrafish Early Embryonic Development by Scanned Light Sheet Microscopy. Science 322, 1065– 1069 (2008).

38. McDole, K. et al. In Toto Imaging and Reconstruction of Post-Implantation Mouse Development at the Single-Cell Level. Cell 175, 859–876.e33 (2018).

39. Biddy, B. A. et al. Single-cell mapping of lineage and identity in direct reprogramming. Nature 564, 219 (2018).

40. Ronneberger, O., Fischer, P. & Brox, T. U-Net: Convolutional Networks for Biomedical Image Segmentation. in Medical Image Computing and Computer-Assisted Intervention – MICCAI 2015 (eds. Navab, N., Hornegger, J., Wells, W. M. & Frangi, A. F.) 234–241 (Springer International Publishing, 2015). doi:10.1007/978-3-319-24574-4_28.

41. Zhang, A. W. et al. Probabilistic cell-type assignment of single-cell RNA-seq for tumor microenvironment profiling. Nat. Methods 16, 1007–1015 (2019).

