## Supplementary materials for "Exploration and analysis of molecularly annotated, 3D models of breast cancer at single-cell resolution using virtual reality"

**SUPPLEMENTARY MATERIAL**

1. **Supplementary Figures and Legends**
2. **Supplementary Figures**
3. **Supplementary Table**
4. **Supplementary Video Legends**

### SUPPLEMENTARY FIGURES AND LEGENDS

**Supplementary Figure S1.** Spatial plots for all IMC channels of the final segmented 3D multi-modal syngeneic 4T1 dataset (after co-registration). Only one 2D section is shown for clarity.

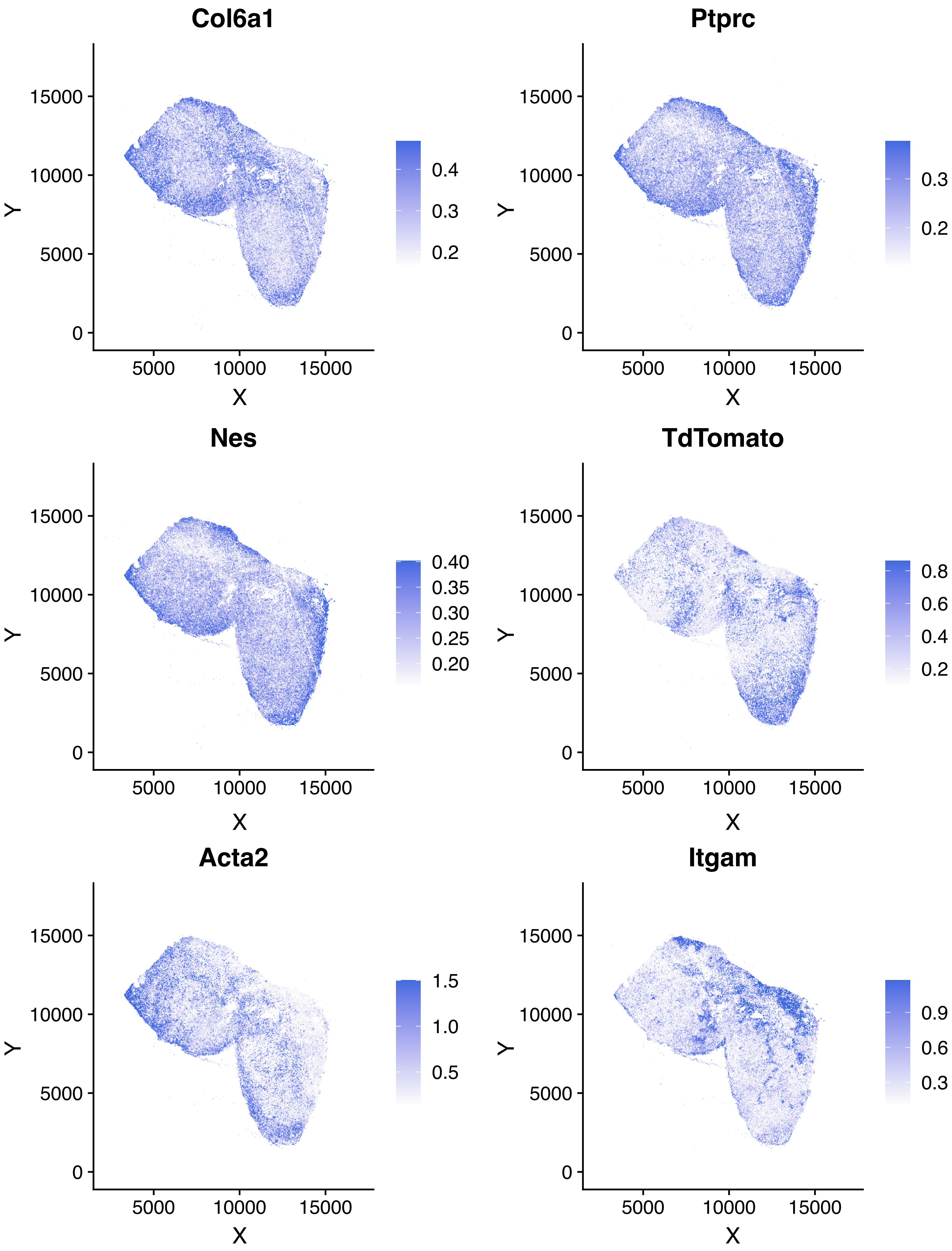
Col6a1
Ptprc
Nes
TdTomato
Acta2
Itgam
X
Y
0
5000
10000
15000
0.2
0.3
0.4
0.20
0.25
0.30
0.35
0.40
0.6
0.8
0.5
1.0
1.5
0.9

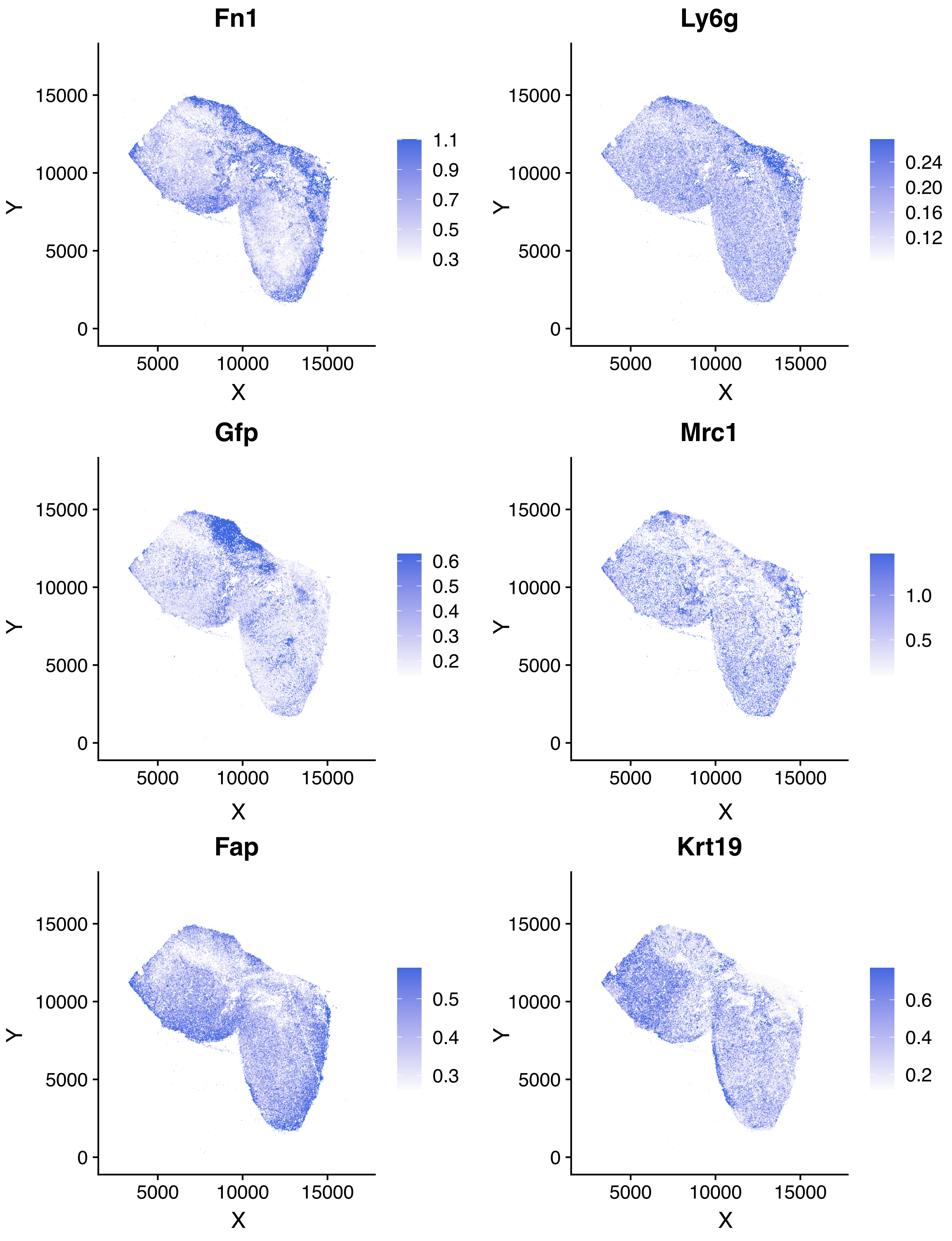
Fn1
Ly6g
Gfp
Mrc1
Fap
Krt19
X
Y
0
5000
10000
15000
1.1
0.9
0.7
0.5
0.3
0.24
0.20
0.16
0.12
0.6
0.4
0.2
1.0

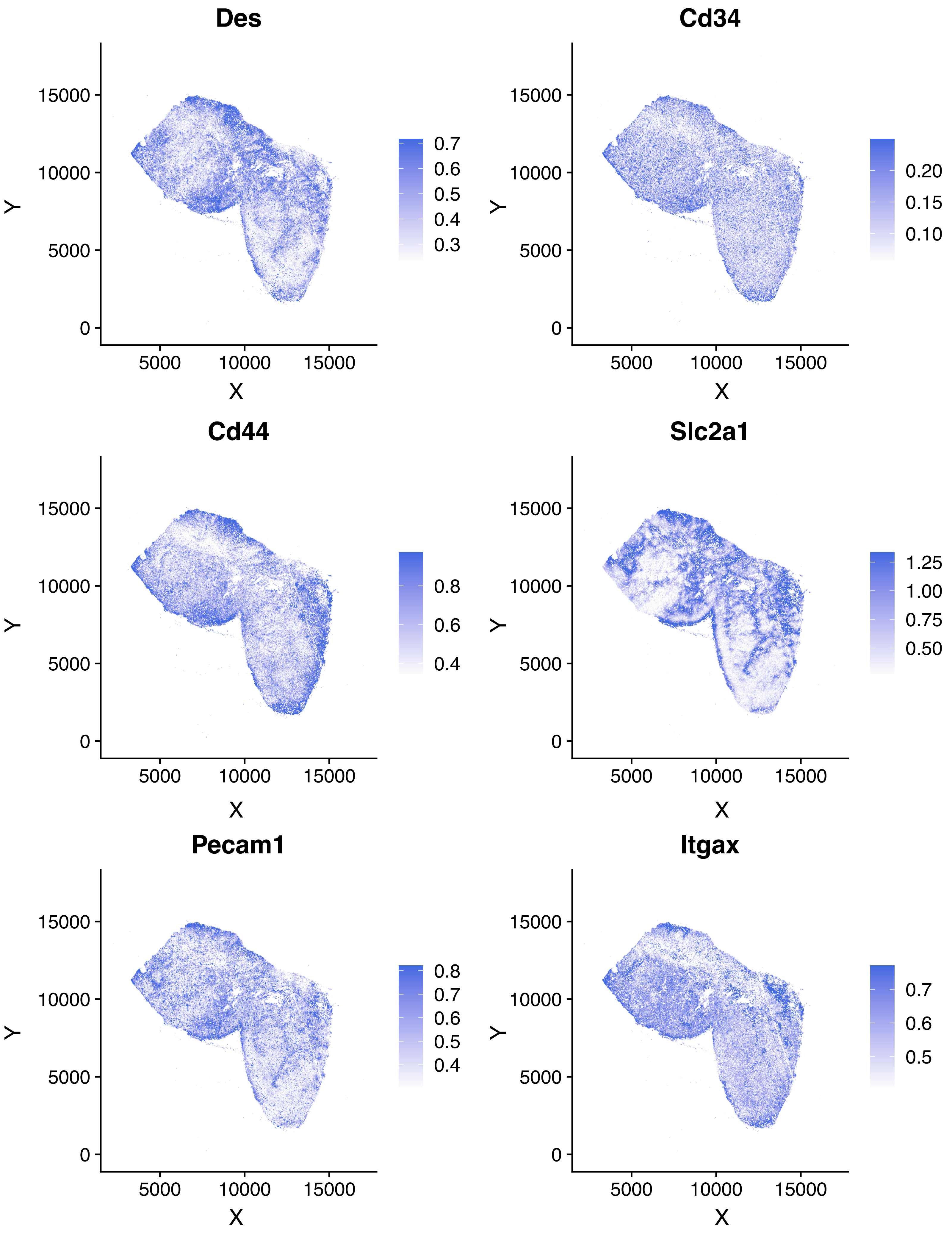
Des
Cd34
Cd44
Slc2a1
Pecam1
Itgax
X
Y
0
5000
10000
15000
0.7
0.6
0.5
0.4
0.3
0.20
0.15
0.10
0.8
0.6
0.4
1.25
1.00
0.75
0.50
0.8
0.7
0.6
0.5
0.4
0.7
0.6
0.5

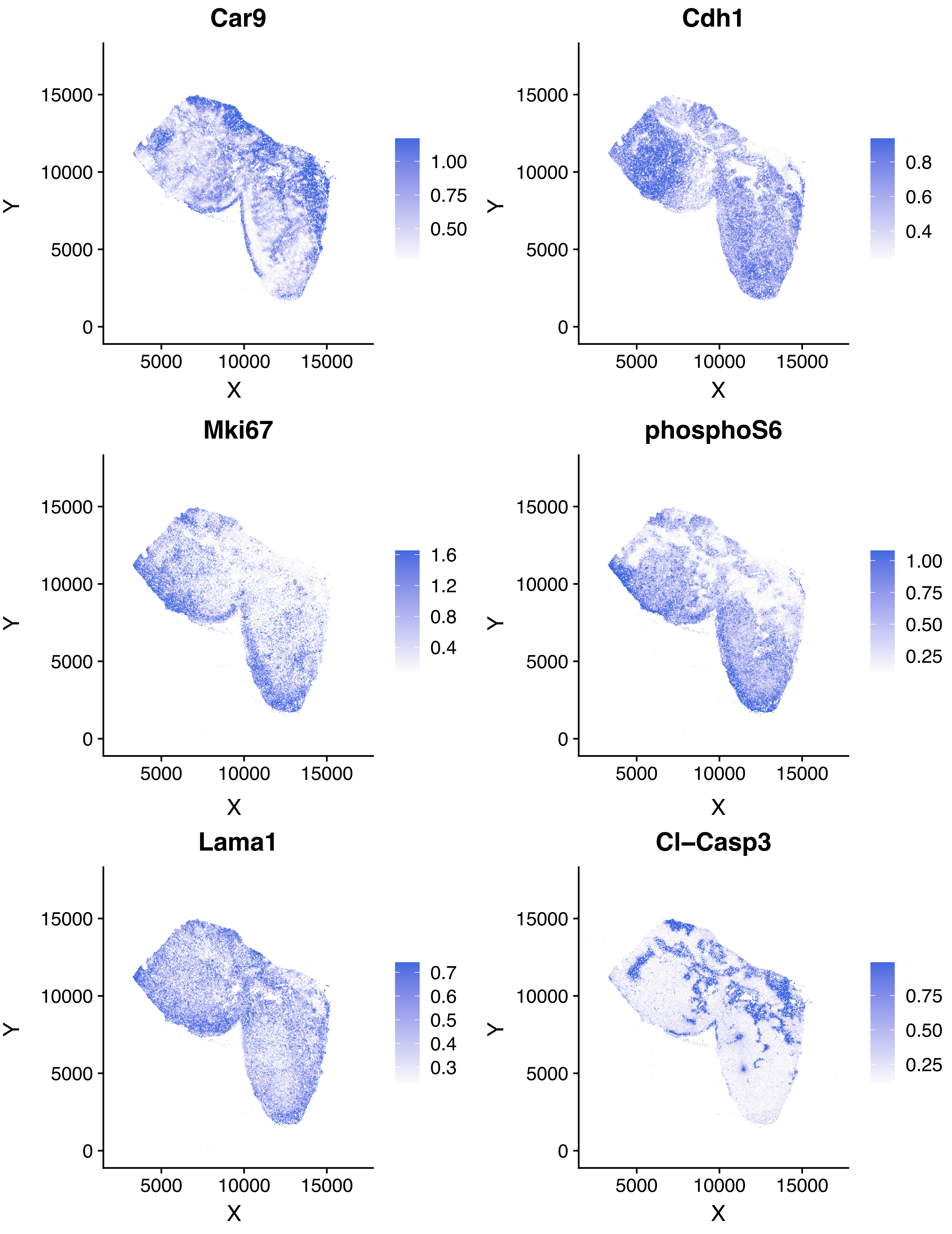
Car9
Cdh1
Mki67
phosphoS6
Lama1
Cl–Casp3
X
Y
0
5000
10000
15000
1.00
0.75
0.50
0.8
0.6
0.4
1.6
1.2
0.8
0.4
1.00
0.75
0.50
0.25
0.7
0.6
0.5
0.4
0.3
0.75
0.50
0.25

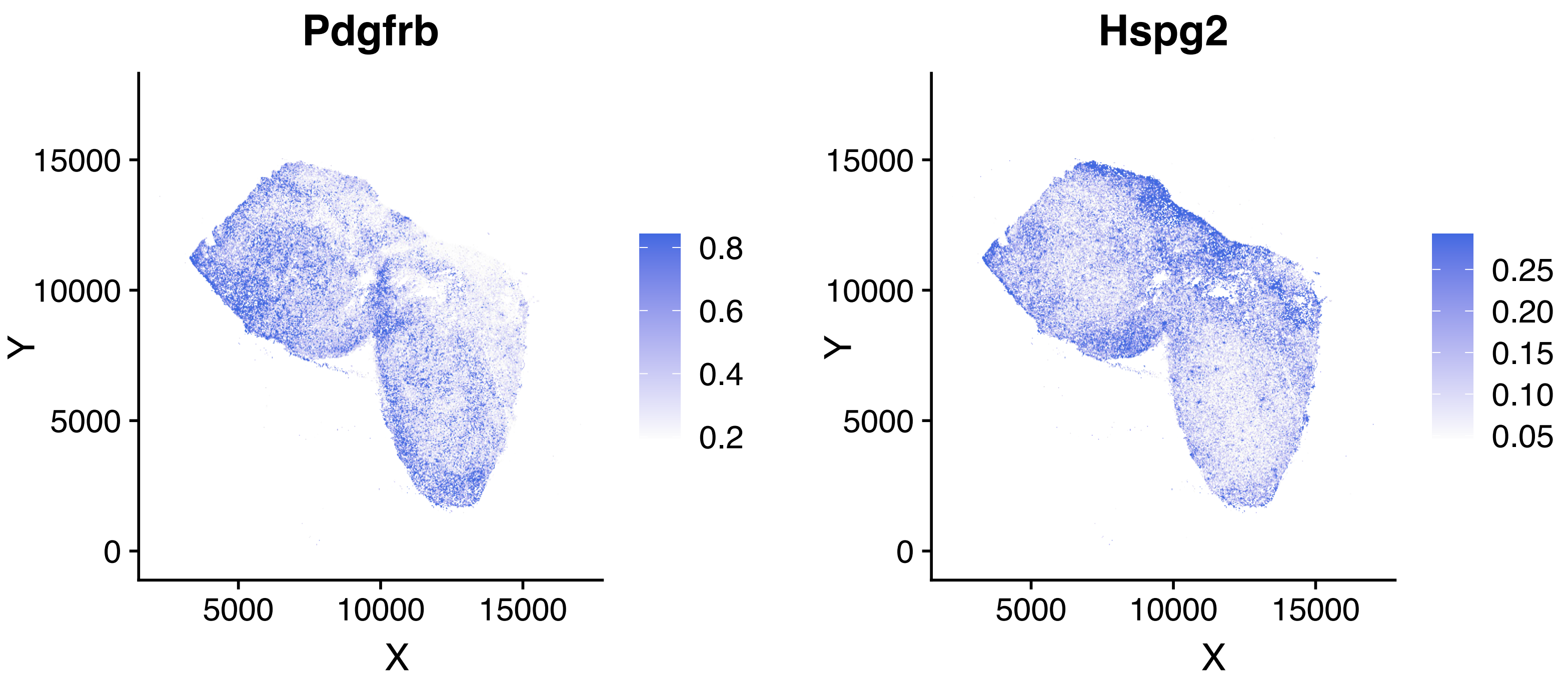
Pdgfrb
Hspg2
15000
10000
5000
0
Y
5000
10000
15000
X
0.8
0.6
0.4
0.2
0.25
0.20
0.15
0.10
0.05

**Supplementary figure S2:** A: Wrist tools in Theia. From top left to right: snapshot camera (currently selected), delete object/tile, teleport, select/deselect cells, pointer / remote manipulation tool, fly tool. B: Contextual help in Theia. Hovering a hand controller over an item of the VR space causes a suggestion text to appear.

A

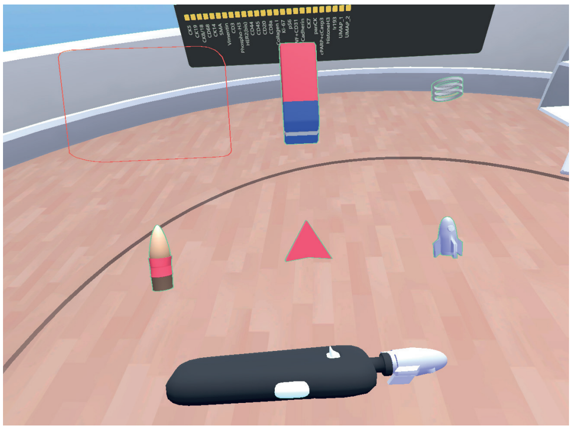

B

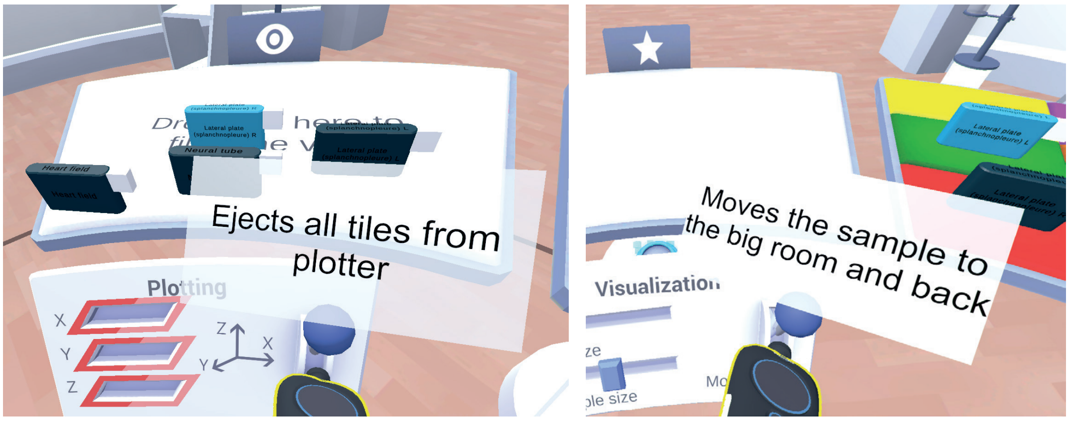


Supplementary Figure 2

### SUPPLEMENTARY TABLES

**Supplementary table 1:** Full list of antibodies, clones, suppliers, catalogue number and concentrations used for the IMC panel.

| Target | Metal | Clone | Source | Catalogue | Reactivity | Working [] |
|---|---|---|---|---|---|---|
| Collagen VI | 141Pr | Polyclonal | Abcam | ab6588 | Human, Mouse | 5 ug/ml |
| CD45 | 142Nd | 30-F11 | Novus | NB100-77417SS | Human, Mouse | 5 ug/ml |
| Vimentin | 143Nd | EPR3776 | Abcam | ab193555 | Human, Mouse | 2.5 ug/ml |
| F4/80 | 144Nd | BM8 | Thermo | 14-4801-82 | Mouse | 10 ug/ml |
| Fsp-1 | 145Nd | Polyclonal | Merck | 07-2274 | Human, Mouse | 5 ug/ml |
| Nestin | 146Sm | Rat 401 (4D4) | Thermo | 14-5843_82 | Human, Mouse | 10 ug/ml |
| TdTomato | 147Sm | Polyclonal | St Johns Lab | STJ140002 | - | 5 ug/ml |
| aSMA | 148Nd | 1A4 | Thermo | 14-9760-82 | Human, Mouse | 5 ug/ml |
| CD11b | 149Sm | EPR1344 | Fluidigm | 3149028D | Human, Mouse | 5 ug/ml |
| Fibronectin | 150Nd | Polyclonal | Novus | NBP1-91258 | Human, Mouse | 10 ug/ml |
| Ly6g | 151Eu | RB6-8C5 | Abcam | ab25377 | Mouse | 5 ug/ml |
| GFP | 152Sm | EPR14104 | Abcam | ab220802 | - | 10 ug/ml |
| MMR/CD206 | 153Eu | Polyclonal | R&D Systems | AF2535 | Mouse | 5 ug/ml |
| FAP1 | 154Sm | Polyclonal | Biorbyt | orb227989 | Human, Mouse | 10 ug/ml |
| Cytokeratin 19 | 155Gd | EPNCIR127B | Abcam | ab133496 | Mouse | 5 ug/ml |
| Desmin | 156Gd | Y66 | Abcam | ab216616 | Human, Mouse | 2.5 ug/ml |
| CD34 | 158Gd | RAM34 | Thermo | 14-0341-82 | Mouse | 5 ug/ml |
| CD44 | 160Gd | IM7 | Thermo | 14-0441-82 | Human, Mouse | 5 ug/ml |
| Glut-1 | 161Dy | EPR3915 | Abcam | Ab196357 | Human, Mouse | 2.5 ug/ml |
| CD31 | 162Dy | EPR17259 | Abcam | ab225883 | Human, Mouse | 5 ug/ml |
| Collagen I | 163Dy | Polyclonal | Novus | NB600-408 | Human, Mouse | 5 ug/ml |
| CD11c | 164Dy | Polyclonal | Biorbyt | orb13554 | Human, Mouse | 10 ug/ml |
| Collagen IV | 165Ho | Polyclonal | Abcam | ab6586 | Mouse; Human | 7.5 ug/ml |
| CA9 | 166Er | Polyclonal | Novus | NB100-417 | Human, Mouse | 5 ug/ml |
| E-Cadherin | 167Er | 36/E-Cadherin | BD | 610182 | Human, Mouse | 5 ug/ml |
| Ki67 | 168Er | B56 | Fluidigm | 3168022D | Human, Mouse | 5 ug/ml |
| pS6 | 170Er | D57.2.2E | Cell Signaling | 4858BF (custom) | Human, Mouse | 2.5 ug/ml |
| Laminin | 171Yb | Polyclonal | Novus | NB300-144 | Human, Mouse | 1 ug/ml |
| Cleaved Caspase | 172Yb | 5A1E | Fluidigm | 3172027D | Human, Mouse | 5 ug/ml |
| CD3 | 173Yb | CD3-12 | Abcam | ab11089 | Human, Mouse | 10 ug/ml |
| PDGFRB | 174Yb | 28E1 | Cell Signaling | Custom order | Human, Mouse | 5 ug/ml |
| Perlecan | 175Lu | A7L6 | Thermo | MA1-06821 | Human, Mouse | 5 ug/ml |
| CD4 | 176Yb | EPR19514 | Abcam | ab221775 | Mouse | 10 ug/ml |

**SUPPLEMENTARY VIDEO LEGENDS**

Supplementary videos can be downloaded from

http://www.hannonlab.org/Bressan_et_al_supplementary/supplementary_videos.zip .

A description of the individual videos is provided below.

**Supplementary video 1. 3D volume produced after mosaicking and re-alignment of the raw STPT images.** Channel 2 (TdTomato+ cells, parental 4T1 line) is in red and channel 3 (GFP+ cells, 4T1-T clone) in green. The whole sample (approximately 1 cm in size) is shown initially, followed by a zoom-in to the level of single cells (individual green/red blobs) to highlight the full resolution of the dataset. The spatial heterogeneity of parental 4T1 / T clone can be easily detected.

**Supplementary video 2. 3D volume produced after co-registration of the IMC datasets to the STPT coordinate space.** Markers for TdTomato, GFP, Ki67 (proliferation), CD31 (endothelial vessels), and Cd11b (myeloid cells) are shown, as well as nuclear counterstain (iridium intercalator). GFP+ tumour cells (4T1-T clone) are spatially separated from proliferative areas and tend to overlap with myeloid infiltration.

**Supplementary video 3. Highlights of the IDC2 3D-IMC model in virtual reality.** First sequence: two sub-populations of basal cells (CCK5hi and SMAhi) are highlighted in red and blue and visualized in 3D. The tubular ductal structures (lined preferentially by one or the other population) can be appreciated much better in VR than using 2D images. Second sequence: co-localization of endothelial cells (defining a blood vessel) and CD8+ T cells. Third sequence: expression pattern of the phospho-S6 (Ser235/236) marker visualized in red. Increased expression of the marker can be seen in the basal cells and the tumour cells proximal to the lining of the invaded ducts.

**Supplementary video 4. Basic manipulation of the multi-modal NSG 4T1 model in virtual reality.** The 3D model is loaded by inserting the corresponding tile into the sample slot of the “projector” tool in the VR environment. Each cell is displayed as a 3D object with shape and colour depending on its cell type. An information panel provides basic information on the sample. Sample rotation (performed by grabbing and moving the projector tool), scaling, cell size adjustment and Z-span adjustment are demonstrated (first person view).

**Supplementary video 5. Basic exploration of the IDC2 3D-IMC model in virtual reality.** Video is shown in third person (the blue avatar corresponds to the user). The model is loaded by moving the tile in the projector tool. Once the sample is loaded, the general structure of the model can be appreciated (tumour-filled mammary ducts, in green, separated by stromal channels in purple and by an immune-infiltrated region around a blood vessel in grey/cyan/red). Movement of the sample, scaling, and cell type adjustment are demonstrated

**Supplementary video 6. Visualization of different cell populations from the multi-modal NSG 4T1 model using virtual reality.** Gfp+ and Tdtomato+ are first displayed in green and red, then stromal cells, myeloid cells (labelled as monocytes) and proliferating cells are displayed in orange, green and red.

**Supplementary video 7. Visualization of different cell populations from the IDC2 3D-IMC model.** Basal $SMA^{hi}$ cells and B cells are visualized in blue and red, then stromal cell and tumour cells are visualized in pink and green. In the end of the video all four cell types are visualized while the sample is scaled.

**Supplementary video 8. Contextual help in Theia.** Hovering a hand over a tool for a few seconds brings up a suggestion text to guide the user towards the more relevant actions to analyse the sample, facilitating a smooth learning curve.

**Supplementary video 9. Cell type cycling in virtual reality for the multi-modal NSG 4T1 model.** Each cell population (as defined in the packaged data) is visualized for a given (and user-definable) time in a looping cycle. (Endothelial cells: red - Hypoxic cells: pink – Myeloid cells: blue - Mrc1+ cells: cyan – Stromal cells: green - Tumour GFP+ cells: yellow - Tumour Tomato+ cells: orange, Tumour Ki67+ proliferating cells: lime) (first person view).

**Supplementary video 10. Cell type cycling for the IDC2 3D-IMC model. (third person view).** Each cell population composing the sample is visualized sequentially in different colours. The populations appearing in order are: CK14-high tumor, Undefined5, Basal CK5-high, CD8+ T cells, Endothelial, B cells, Undefined4, Basal SMA-high, Macrophage, general tumour cluster, CK19-high tumour, Ki67-high tumour, CD44-high tumour, Undefined2, Undefined1, CD8- T cells, Stroma, Luminal tumour, Undefined3, CK8/18-high tumour, CK14-high tumour, Undefined5. Distinct spatial patterns can be seen for most cells, including heterogeneity in the population of tumour cells filling different ducts (i.e. $CK19^{hi}$ tumour cells) (third person view).

**Supplementary video 11. Cell selection by marker intensity**. The SMA marker was used to define a $SMA^{hi}$ cell population directly from within the VR environment. Adjusting the threshold expressing selects only stromal and basal cells (pink and blue)

**Supplementary video 12. marker intensity colour mapping.** The Ki67 marker is displayed in red in the IDC2 model. Gene intensity is mapped to colour level and the min-max levels can be adjusted freely. Higher expression can be seen inside the tumour-filled ducts.

**Supplementary video 13. Sample and population information panels.** The population information panel includes a marker expression heatmap which is dynamically updated as new cells are selected from the sample using the selector tool (here demonstrated using the IDC2 3D-IMC sample)

**Supplementary video 14. VR fly-in of the multi-modal NSG 4T1 model.** By default the 3D models visualized in Theia are rendered to a size roughly matching that of the user, and can be manipulated by grabbing the "projector" tool below them. However, it is possible to increase the model size many-fold and to fly inside it to identify individual structures of cells. This operation is demonstrated here for the multi-modal NSG 4T1 model. All cells in the model are visualized in grey. GFP+ cells (4T1-T clone) are in red. (first person view).

**Supplementary video 15. VR fly-in of the IDC2 3D-IMC model.** Fly-in exploration of the IDC2 model. Stromal cells are in pink, tumour cells in green, basal $SMA^{hi}$ cells in blue, B cells in red (third person view). Even though the model is completely filled with cells, adjusting the sample and cell size can produce some empty space between cells which allows appreciation of individual cells. Tissue heterogeneity can be immediately perceived using both colours and cell shapes.

**Supplementary video 16. Example of the marker 3D plot tool.** The SMA, Ki67 and CD68 markers were used a produce a 3D plot for the IDC2 3D-IMC model by inserting the corresponding tiles inside the slots of a "plotting" tool, each corresponding to an axis of the plot. The min/max of each axis can be modified by accessing a label on each gene tile. In this video, the scale of the SMA marker is modified, resulting in $SMA^{hi}$ stromal and basal cells becoming more separated from the rest of the cells in the model.

**Supplementary video 17. Integrated dimensional reduction tool.** Cells from the IDC2 model are selected using the cell selection tool in Theia, and the dimensional reduction tool is used to calculate a UMAP embedding on the fly. Other dimensional reduction strategies can be enabled by manipulating the python script attached to the VR tool.

**Supplementary video 18. Example of round-trip analysis on th IDC2 3D-IMC sample.** A sub-population of cells is highlighted in the UMAP dimensional reduction plot using the cell selection tool, saved as a new tile, coloured in red, and re-mapped to spatial coordinates, revealing a clustered spatial pattern.

**Supplementary video 19. Differential expression tool.** Differential expression is calculated using the integrated differential expression tool in Theia. In this example, the two basal cell populations ($SMA^{hi}$ and $CK5^{hi}$) from the IDC2 3D-IMC model are selected and compared, identifying Vimentin and phospho-S6 (Ser235/236) as significantly enriched markers in the $CK5^{hi}$ basal cell population. Adjusted p values are calculated through the t-test implementation of the *scanpy* package.

**Supplementary video 20. Neighbourhood search.** A custom population is defined from the IDC2 3D-IMC model using the cell selector tool, and expanded using the "Select nearby neighbours" function. Distance is expressed as a function of the model size (each distance unit corresponds to approximately 1/2000$^{th}$ of the maximum model dimension). The initially selected population is shown in red and the neighbouring one in green

**Supplementary video 21. Example of multi-user exploration of datasets using Theia**. Three users are interacting with the IDC1 3D-IMC sample, selecting different cell populations/types and manipulating the model. Both heads (coloured spheres) and hands (black controller shapes) are tracked for each user.

**Supplementary video 22. Exploration of a non-spatial single-cell dataset using Theia.** VR can be used for traditional single-cell datasets by visualizing 3D dimensional reduction embeddings as coordinates. In this case scRNAseq data produced from a 4T1 tumour produced in a NSG mouse is visualized (any other dataset can be converted via a provided python script). Two populations of cells including stromal or tumour cells are selected and a differential expression is calculated on the fly. The stromal cells show upregulation of *Sparc,* (osteonectin), *Bgn* (biglycan) and collagen genes, consistently with its identification.

**Supplementary video 23. Visualization of the mouse embryo development dataset from McDole et al. in virtual reality.** Segmented cells were visualized and the embryo shape can be seen evolving over time. Time starts at embryonic day 6.5 (early streak stage) and ends at embryonic day 8.5 (somites stage). Initially each cell lineage is mapped to a separate colour, with most cells appearing red (the colour of "undefined" lineages, particularly abundant in the early time points) then the neural tube is selected in blue, and finally the neural tube and heart field are selected in green and red, respectively. The elongation of the neural tube over time can be easily detected, as well as many other features of embryo development.

**Supplementary video 24. Theia volumetric viewer.** A 3D image produced by STPT of a mouse mammary gland (NSG-GFP+ strain – see methods). GFP+ ductal structures can be seen in green.

Second-harmonic generation (SHG) signal from collagen fibres is shown in red. Most ducts are green only, but some structures (larger ducts, i.e. towards the centre, and likely blood vessels) are sheathed in collagen fibres. Manipulation of the voxel rendering (rotation/translation) and use of the cross-section tool are demonstrated.
